# Scalable and systematic hierarchical virus taxonomy with vConTACT3

**DOI:** 10.1101/2025.11.06.686974

**Authors:** Benjamin Bolduc, Olivier Zablocki, Dann Turner, Ho Bin Jang, Jiarong Guo, Evelien M. Adriaenssens, Bas E. Dutilh, Matthew B. Sullivan

**Affiliations:** Department of Microbiology, Ohio State University, Columbus, OH, USA; Center of Microbiome Science, Ohio State University, Columbus, OH, USA; EMERGE Biology Integration Institute, Columbus, OH, USA; School of Applied Sciences, College of Health, Science and Society, University of the West of England, Bristol, UK; Center for Study of Emerging and Re-emerging Viruses, Korea Virus Research Institute, Institute for Basic Science (IBS), Daejeon, Republic of Korea; Quadram Institute Bioscience, Norwich Research Park, Norwich, UK; Institute of Biodiversity, Faculty of Biological Sciences, Cluster of Excellence Balance of the Microverse, Friedrich Schiller University Jena, 07743 Jena, Germany; Theoretical Biology and Bioinformatics, Science4Life, Utrecht University, Padualaan 8, Utrecht 3584 CH, The Netherlands; Department of Civil, Environmental and Geodetic Engineering, Ohio State University, Columbus, OH, USA

## Abstract

Viruses are key players in diverse ecosystems, but studying their impacts is technically and taxonomically challenging. Taxonomic complexities derive from undersampling, diverse DNA and RNA genomes with multiple evolutionary origins, and lack of a universal barcode gene. While virus ecogenomics has expanded access to and understanding of the virosphere, available classification tools poorly scale to modern discovery-based datasets, lack taxonomic resolution, and/or are unable to classify novel sequence space. Here we develop, benchmark, and release vConTACT3, a machine learning-based tool that improves scalability and accuracy, adds extensive user-requested features, expands classification to both eukaryote and prokaryote viruses for 4/6 officially recognized realms, and establishes accurate hierarchical taxonomy from genus to order. Application to 48,069 public virus genomes provided new taxonomy assignments for thousands of taxa, revealed support for fewer taxonomic ranks than currently available, and systematically identified taxonomically problematic areas across the virosphere.

## Main

Viruses are increasingly recognized as foundational ecological and evolutionary players in diverse free-living (e.g. oceans^1–3^ and soils^4,5^) and host-associated (e.g., plants^6^, ruminants^7,8^, and humans^9,10^) ecosystems. Problematically, diverse virus lifestyles and myriad possible genomic structures (i.e. single or double-stranded, DNA or RNA) make formalizing taxonomic classification challenging, with obstacles remaining as follows. *First and foremost*, differences in evolutionary rates across virus sequence space (the virosphere) preclude single demarcation criteria across all realms, resulting in a patchwork of approaches for currently approved taxa by the International Committee on the Taxonomy of Viruses (ICTV)^11–13^. This makes comparing analogous taxonomic ranks across vastly different virus lineages challenging. *Second*, the ICTV recently made available 15 taxonomic ranks^14^ (species to realm), but thus far none of the 14,690 ICTV-annotated virus species have ranks assigned, with “sub-” rank assignments (subrealm, subkingdom, subphylum, etc.) optional and not in use for two-thirds of these taxa. Further, no global survey of the sampled virosphere has yet evaluated to what extent the data support the need for a 15-rank structure. *Third*, virus taxonomy is not static, but an ongoing effort to reflect the available data at a given time. As most of the 10^31^ viruses on the planet have not been sequenced, current taxonomic descriptions are derived from a fraction of Earth’s virosphere. For example, even the most extensive virus genomic resource (IMG/VR^15^) harbors only ~15.3 million virus genome fragments, which is many orders of magnitude fewer than the number of viruses that exist on the planet, and ICTV classification is only available for less than 0.01% of the IMG/VR sequences. Though it is unknown how many virus ‘types’ the 10^31^ virus particles will represent, the fact that virus surveys, particularly in newly studied ecosystems, routinely identify novel viruses that are not taxonomically classifiable at low ranks suggests that we have a long way to go to capture the many forms of virus genomes that exist on Earth.

Currently, while there is growing ICTV agreement that a genome-based, evolutionary framework is needed to support a universal virus taxonomy^16–19^, no consensus tool or platform exists to implement this vision. Several tools exist that can assign new sequences to *known* taxa, but because those tools lack an underlying ruleset or statistical framework they cannot also cannot create new taxa where needed. Examples include tools that utilize ‘hallmark genes’ (genes shared among a group of viruses, but not universally across the virosphere)^20–22^ or their translated proteins to align and analyze for genome-wide shared gene content (VirClust^23^), HMM profiles or protein families (GRAViTy^24^, VPF-Class^25^, geNomad^26^), or genomic signals (e.g. VIRIDIC^27^, PASC^28^). Recent developments combining hierarchical clustering with core protein or gene marker detection provide annotation and near-reference taxonomy (VirClust^23^, Cenote-Taker^29^), but they are limited in scalability and cannot create new taxa. Among these, only geNomad is scalable as it leverages a large marker gene dataset, with demonstrated assignment accuracy to the family level for near-reference genomes^26^, but higher-level taxa are problematic and it cannot create new taxa.

To date, the only approach that is both scalable and able to create new taxa uses gene-sharing networks to identify statistically-supported ‘virus clusters^‘30^ (VCs) in sequence space that are benchmarked against ICTV taxonomy. This method has been extensively benchmarked for dsDNA phages and at a single rank (genus), and the capability has been formalized into tools (vConTACT^31^, vConTACT2^32^) that have been crucial in classifying these phages in both detailed taxonomic as large-scale metagenomic studies^33^. However, even vConTACT2 has limitations that significantly hamper viral discovery.

Here we introduce vConTACT3, which significantly expands upon the above. Specifically, vConTACT3 better scales, provides hierarchical classification across 5 ranks from genus to order, and does so beyond just dsDNA prokaryotic viruses (**Fig. 1**). Further, data exploration is enhanced via six user-requested outputs (Supplemental Note 1), and an overhauled reference database that allows run-to-run comparisons. Together, these improvements bring us much closer to implementing the ICTV vision of a genome-based universal virus taxonomy^18^.

**Figure 1.**
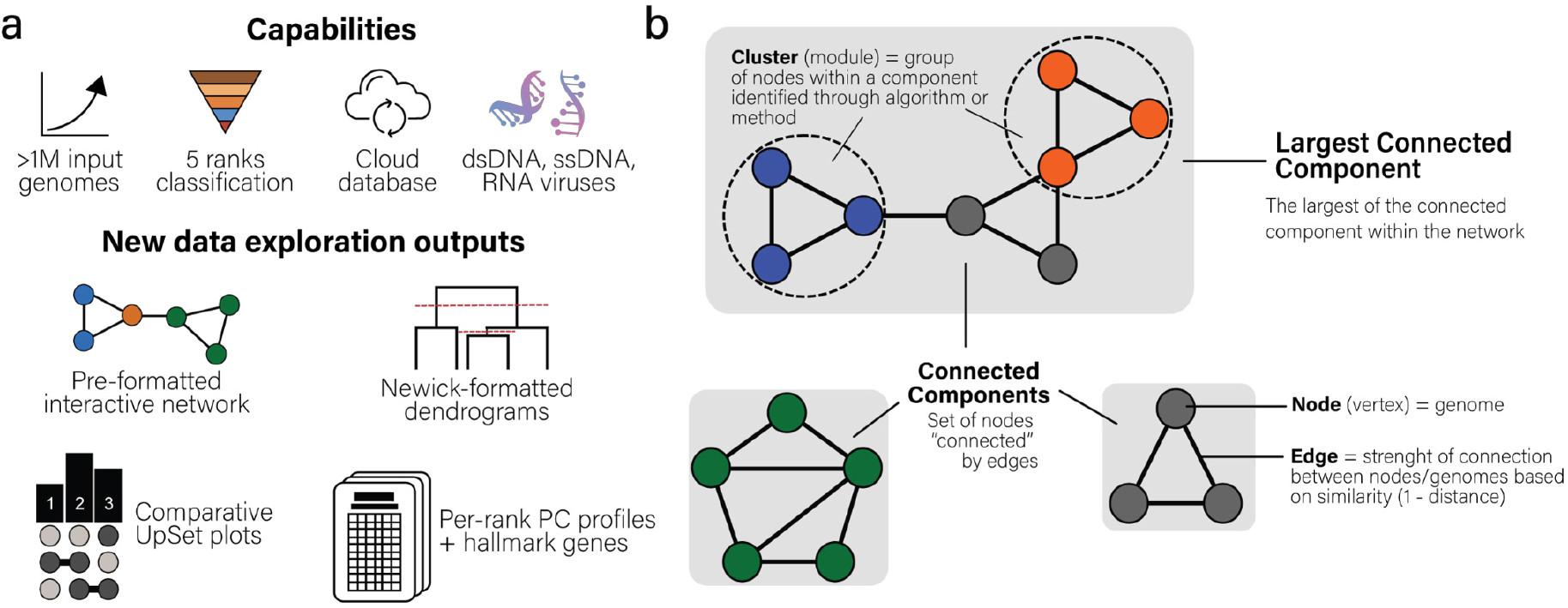
vConTACT v.3.0 new features overview. (**a**) An overview of features between the three vConTACT family of tools. (**b**) A conceptual overview of a network and its parts. Networks (known as graphs) are fundamentally composed of a node (also known as a vertex), connected to another node through an edge. Nodes in vConTACT3 are genomes, whereas edges in vConTACT3 are defined by the number of genes shared and conversion into a between-genomes similarity score. Each network is separated into connected components, with the largest known as the “largest connected component”. Within these connected components are clusters (highlighted by blue and orange) that are ideally defined through an algorithm-aided formal statistical approach.

## Results and Discussion

### vConTACT3 expands classification ranks and realms

Our first challenge was to expand upon single-rank genus-level assignments, while maintaining high accuracy. Prior vConTACT versions^31,32^ produced VCs based on shared-gene content that approximated genus-rank groupings, as benchmarked against predominantly tailed dsDNA phages that belonged to the class *Caudoviricetes* (formerly order *Caudovirales*)^34^. Classification was limited to genus-rank because (i) thresholds for genus ranks were defined whereas the 15-rank formalization came later^14^, and (ii) vConTACT2’s VCs, produced by the ClusterONE algorithm struggled with accommodation and/or accuracy at lower and higher ranks.

To counter the difficulty with vConTACT 2, we completely redesigned the vConTACT pipeline. Specifically, we removed ClusterONE (which produced genus-rank clusters) and expanded clusters to extend to the largest connected component(s) (‘LCC’) in the network, optimizing these networks to maximize their taxonomic concordance with ICTV, enabling hierarchical clustering spanning all genomes within the LCC. Only the LCC was used for benchmarking because it consisted of the greatest realm membership, and optimization against all connected components (CCs) would introduce multiple potential distances per realm and rank, and we note that those may differ in smaller CCs. Distances were optimized to fit the revised ICTV taxonomy, allowing for distinct thresholds in different lineages to resolve uneven rates of evolution, and potential multi-realm LCCs drawn together via horizontal gene transfer were resolved as follows.

We first empirically determined optimal distances to best fit existing, hierarchically ranked taxonomic assignments for extant viruses in NCBI RefSeq database (~20K genomes) that spanned 6 realms and 3 host domains. To do this, we evaluated >60 million parameter combinations via iterative and incremental clustering of protein clusters (PCs) PCs at various clustering identities. This reference dataset of 6 realms and 3 host domains and the parameter combinations analyzed both go well beyond what we^31,32^ or others^35^ have tested previously since most tools optimized on select virus realms and/or ranks. In contrast, for vConTACT3 we chose to optimize, benchmark, and assess performance across 6 realms (*Ribozyviria, Adnaviria, Monodnaviria, Riboviria, Duplodnaviria, Varidnaviria*) and all 3 host domains (*Bacteria, Archaea, Eukarya*) – and did so across 8 virus taxonomic ranks (realm, kingdom, phylum, class, order, family, subfamily, genus). Analyses were evaluated using 4 distance metrics, 8 protein clustering identities, 100 pairwise distance cutoffs, and 25 different values of minimum genes shared (**see Methods**). In all, analyses of these >60 million evaluated combinations comprehensively optimized realm- and rank-specific cutoffs within existing ICTV taxonomy (summarized in **Fig. 2**). Since only the LCC was used to optimize distances, discrepancies in accuracies for each realm and rank may arise principally due to disparate distances between the smaller CCs and LCC between viruses of the same realm.

**Figure 2.**
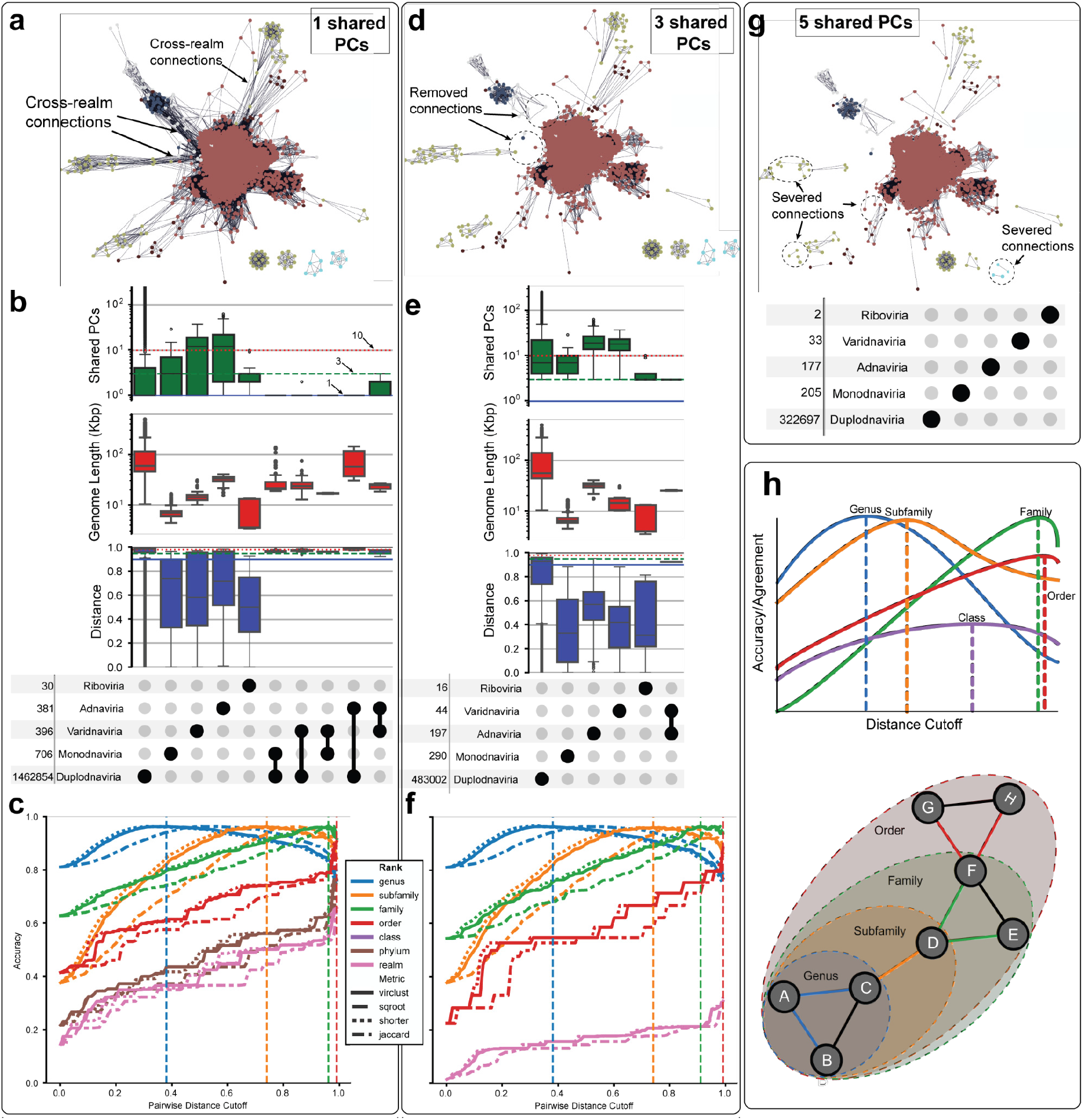
Benchmarking vConTACT3 with 60 million combinations. (**a**) Network with nodes (i.e. genomes) colored by realm, with edges representing a minimum of 1 shared gene between each genome. Annotations indicate cross-realm connections (when two genomes of different realms are connected) (**b**) UpSet plots of realms, derived from the network in (a), showing within and between realm edges. Top-most plot of the UpSet shows counts of *shared PCs*, with horizontal lines indicating 1, 3, and 10 shared PCs (dashed black, green and red) lines, *genome lengths*, and *distances*, with horizontal lines indicating 0.90, 0.95 and 0.98 (solid blue, dashed green, dotted red) lines. (**c**) Line plots of accuracies at distances from 0 - 0.99, colored by taxonomic rank. Dashed lines represent the distance metric type (Jaccard, “SqRoot”, ‘VirClust” and “Shorter”) employed. (**d-f**) Corresponding network, UpSet plot, and accuracy plot of A-C, but requiring a minimum of 3 shared genes to establish a relationship between genomes. Dashed circles in (d) highlight some instances of where increasing the minimum shared genes disrupted/severed edges between previously cross-realm connections. UpSet plot shows only 3 edges/PCs shared between viruses of the *Varidnaviria* and *Adnaviria*. (**g**) Networks as in (a) and (d), but with 5 minimum shared PCs. Annotations indicate severed connections/edges *between genomes of the same realm*. (**h**) Representative network showing correspondence between maximal accuracy and rank selection for each clustering identity (0.3 - 0.7) and minimum shared gene (1 - n). Distance cutoff determined in (c) and (f) is applied to the network(s) prior to hierarchical clustering.

Together, these analyses revealed that optimized cut-off values varied by both the host domain and realm, reflecting the different taxonomic methods and cutoffs used by the ICTV to handle the highly varied genomes across the virosphere. *First*, protein clustering identities (used to establish shared protein clusters) in the prokaryote-infecting *Duplodnaviria* tended to increase from order (30% sequence identity) to genus (70%), with decreasing pairwise distance cutoffs (0.99 to 0.55 of the similarity score) (30% identity **Fig. 2c, 2f;** all identities **Extended Data Fig. 2**), whereas those from eukaryote-infecting viruses used only two clustering identities (30% and 40%), and cutoffs were narrower (0.99 to 0.74) (**Extended Data Fig. 3**). This meant that to maximize ICTV agreement across ranks, 4 protein clustering identities and greater range of pairwise distances were required for prokaryote-infecting viruses compared to only 2 protein clustering identities for eukaryote-infecting viruses.

Further, two realms – *Adnaviria* and *Varidnaviria –* only needed one (30%) protein clustering identity whether viruses infected prokaryotes (**Extended Data Fig. 4**) or eukaryotes (**Extended Data Fig. 5**). *Second*, prokaryote-infecting virus thresholds were consistently shifted lower/higher than those of eukaryote-infecting viruses (**Extended Data Fig. 6**). For example, across *Duplodnaviria* the genus, subfamily and family of those infecting eukaryotes correspond to subfamily, family, and order of those infecting prokaryotes. A similar shift is seen for *Adnaviria* and *Varidnaviria*, where the genus ranks agree, but prokaryote-infecting viruses lack a subfamily rank and the cutoff of the prokaryote-infecting family rank corresponds to that of subfamily in eukaryote-infecting viruses. *Third*, these evaluations also revealed that 1-5 minimum *shared* genes provided the greatest discriminatory power, because any more than that limited higher taxonomic ranks (beyond family) being assigned (**Fig. 2a, 2d, 2g**). Together these results show that the virus genomic sequence space can be related to ICTV taxonomy through tuning protein similarities and group-specific threshold selections.

Beyond parameter tuning to optimize ICTV agreement, the second hurdle encountered was to maintain separation and accuracy within potential multi-realm LCCs that are drawn together due to horizontal gene transfer, convergent evolution, and/or homoplasy^36^. For example, we found that a naive gene-sharing network using the above optimized distances without shared gene minimums worked well most of the time, but could also produce problematic findings – one LCC connected 3 different realms (*Duplodnaviria, Varidnaviria* and *Monodnaviria*) and two other LCCs connected 2 realms (both *Monodnaviria*-*Varidnaviria*) (**Fig. 2a**). While rare (~0.03% of the total connections; **Fig. 2b**) and resolvable via additional hierarchical decomposition, we sought a globally-relevant threshold-based cutoff to enable their systematic resolution.

To explore this, we pre-filtered the gene-sharing networks by removing poorly supported edges between genomes (2 or fewer shared genes) and this largely eliminated the problem as 99.99% of the “false positive” realm-spanning edges were no longer present (**Fig. 2d, 2e**). Problematically, however, it also removed biologically-accurate genomic connections between virus groups. Specifically, it (i) reduced the sensitivity of hierarchical clustering and/or (ii) removed afflicted genome(s) from clusters, particularly for small genomes like *Ribovaria* and *Monodnaviria*. To mitigate this, we added back genomes that shared 1-2 genes disconnected by pre-filtering where they likely represented deeply-rooted marker genes. Indeed, this resulted in the near-complete removal of all cross-realm connections, while retaining informative within-realm edges. The sole exception was a connection between Adnaviria and Varidnaviria, likely resulting from host-relevant genes (Supplemental Note 2).

Finally, given a much more systematic framework for evaluating virus sequence space, we assessed how many taxonomic ranks might be supported by gene-sharing analyses of virus genomes available in NCBI RefSeq v220 (21,080 genomes). We did this as we noticed that vConTACT3 and ICTV concordance analysis across 3 domains and 6 virus realms revealed that some ranks were accurately distinguished (genus, subfamily, family, and order), but others were not (class, phylum, and kingdom) (**Extended Data Fig. 2-7**). This revealed (**Fig. 2**) that for most realms, only four ranks (genus, subfamily, family, order) could be confidently defined using our approach. This can be seen in the accuracy curve, which peaked at the maximum distance cutoff of 1.0 (i.e., the value for two genomes sharing *no* genes) and reflects viruses from different orders rarely shared genes. This finding was prominent at higher ranks (**Fig. 2c, 2f**) and suggests that kingdom, phylum and class will require information beyond gene-sharing such as clade-specific hallmark genes and/or protein fold data. While these ideas have been discussed elsewhere^18,35^, they have not been globally evaluated as done here. In spite of these limitations, vConTACT3 greatly expands upon single-rank genus-level assignments, to now also assign subfamily, family, order and realm ranks. Finally, we note that the lower rank of species is beyond vConTACT3’s resolution and best captured via sequence similarity of the whole genome (e.g., VICTOR, VIRIDIC or ViPTree, which are commonly used by the ICTV to evaluate new taxonomic proposals^24,37^), or, where sequencing depth allows, more formal population genetic approaches^38,39^.

### vConTACT3 shows concordance with ICTV, enables classification of novel viruses and can inform taxonomic revisions

We next sought to tackle automating rank assignments across taxa, which, due to inconsistent genome similarity thresholds for a given rank, is challenging for classifiers^40^. To this end, we evaluated vConTACT3’s hierarchical rank assignment accuracy against 11,318 prokaryotic viruses in NCBI’s curated RefSeq (v220, accessed Nov. 2023). This revealed that for prokaryotic viruses of *Duplornaviria, Monodnaviria, Adnaviria* and *Varidnaviria*, vConTACT3 averaged 97.6%, 98.7%, 100%, 100%, 90.6%, 90.6% agreement at the ranks of genus, subfamily, family, order, class, and phylum, respectively. This performance by vConTACT3 was almost always (only exception being *Monodnaviria*) better than all available tools that scale, including vConTACT2, VPF-Class, and Genomad, and – critically – also provided these accurate assignments at many more hierarchical ranks (**Table 1**).

**Table 1.** Taxonomic assignment accuracies across tools and realms. The taxonomic predictions from each available scalable tool (VPF-Class, geNomad, vConTACT2, vConTACT3) for sequences across 5 realms were compared against the taxonomies available from the International Committee for the Taxonomy of Viruses (ICTV). Blank spaces represent taxon ranks where the tool is not capable of making a prediction, and accuracies <100% can represent either the tool being inaccurate or the non-systematic nature of ICTV taxonomies in some parts of the virosphere. NP: Not possible.

|  | <u>Adnaviria</u> |  |  |  |  | <u>Varidnaviria</u> |  |  |  |  | <u>Monodnaviria</u> |  |  |  |  |
| --- | --- | --- | --- | --- | --- | --- | --- | --- | --- | --- | --- | --- | --- | --- | --- |
|  | genus | family | order | class | phylum | genus | family | order | class | phylum | genus | family | order | class | phylum |
| Genomad | NP | 91.3% | 100.0% | 100.0% | 100.0% | NP | 83.5% | 96.8% | 74.7% | 74.7% | NP | 92.6% | 99.7% | 99.7% | 99.7% |
| VPF-Class | 93.5% | NP | NP | NP | NP | 90.0% | NP | NP | NP | NP | 91.9% | NP | NP | NP | NP |
| vConTACT2 | 96.5% | NP | NP | NP | NP | 100.0% | NP | NP | NP | NP | 93.5% | NP | NP | NP | NP |
| vConTACT3 | 93.5% | 100.0% | 100.0% | 100.0% | 100.0% | 100.0% | 100.0% | 100.0% | 78.5% | 78.5% | NP | NP | NP | 78.0% | 78.0% |

**Table 1. Taxonomic assignment accuracies across tools and realms.**
|  | <u>Riboviria</u> |  |  |  |  | <u>Duplodnaviria</u> |  |  |  |  |  |
| --- | --- | --- | --- | --- | --- | --- | --- | --- | --- | --- | --- |
|  | genus | family | order | class | phylum | genus | subfamily | family | order | class | phylum |
| Genomad | NP | 86.6% | 100.0% | 100.0% | 100.0% | NP | NP | 95.7% | 92.5% | 100.0% | 100.0% |
| VPF-Class | 91.3% | NP | NP | NP | NP | 80.8% | NP | NP | NP | NP | NP |
| vConTACT2 | 91.3% | NP | NP | NP | NP | 95.9% | NP | NP | NP | NP | NP |
| vConTACT3 | NP | NP | NP | 100.0% | 100.0% | 98.5% | 97.3% | 100.0% | 100.0% | 100.0% | 100.0% |

While promising, recent ICTV reorganization and rank abolishments left some under-assigned ranks, particularly problematic at the order rank where only 20% (1,041 of 5,794) of ICTV bacterial viruses have assignments^34^. To assess vConTACT3’s assignment capability, we applied it to 23,227 prokaryotic virus genomes from the INPHARED database^41^ (release 1 June 2024) and compared assignments against ICTV’s for the subset (4,827) also present in the ICTV Metadata Resource (master species list 39 version 1; https://ictv.global/vmr) for realms *Duplornaviria, Adnaviria* and *Varidnaviria*. Where rank assignment comparisons could be made, we found strong agreement at the order (95.6%) and family (95.7%) ranks. However, subfamilies had lower agreement (only 62%) and examination of ICTV assignments revealed substantial variation in the minimum percentage of shared PCs. The discordance identified at subfamily or family ranks represents areas where manual curation^42^ in the absence of formally defined thresholds was used to make assignments based on the topology of phylogenetic trees. Thus, rather than a fault of vConTACT3, we view discordance at these ranks as areas where vConTACT3 helps identify opportunities for more systematic classification and perhaps even automation as manual evaluation is conducted.

Because genus-level assignments offered the most available manual inspection effort, we explored these assignments in depth. We found that vConTACT3’s automated approach could assign classifications for 97% (4,810) of the 4,827 genomes. Of these, 3,866 (80.1%) fully agreed with existing ICTV assignments, 387 (8%) did not agree, 130 (3%) were discordant with ICTV BVS demarcation criteria and may require re-evaluation, and 424 (9%) represented “edge” cases. These edge cases are where vConTACT3 merged, split or moved species or genera where (i) the inter-genus ANI between genomes was ≥65%, but less than ICTV’s standard of 70%, or (ii) >70% was achieved for some, but not all genomes between two genera. Examples of these rare areas of discordance include virus groups whose genus-level rank assignments were made by the ICTV using phenotype (fuselloviruses^43^) or evolutionary evidence (bacterial microviruses^44^).

Looking at the larger INPHARED prokaryotic virus dataset available, we noted that ICTV assignments were missing at many ranks – 64.5%, 64.5%, 75.7%, and 84.6% were unclassified at the genus, subfamily, family, and order ranks, respectively. To assess vConTACT3 automation, we compared its assignments against manual assessments made via core-genome phylogenies. Specifically, for eight families that included well-curated ICTV virus groups (e.g. the families *Straboviridae, Kyanoviridae, Herelleviridae* and the Felix O1-like viruses), we found vConTACT3 assigned three *novel* orders of *Caudoviricetes* that represented 10% of the available prokaryotic virus genomes in the INPHARED database (2,422 of 23,277 sequences). After extensive manual curation (Supplemental Figure 1), this revealed that vConTACT3 could quickly provide assignments that were in strong agreement with manual pan-genome and core-genome phylogenetic approaches – and does so with virtually no effort on the researchers’ part as compared to the substantial effort required for manual curation.

Given vConTACT3’s strong performance against the 2,422 manually assessed sequences, we next applied it to the full INPHARED dataset. This resulted, automatically, in floods of new assignments including 3,113 new genera (with 964 existing references and merging 72 others), 1,335 new subfamilies (83 existing and 12 merged), 803 new families (with 67 existing) and 192 new orders (with 9 existing). Even when excluding singleton taxon ranks (i.e., ranks where only a single genome representative is available), these new assignments were still many, with 728 new genera, 697 new subfamilies, 493 new families, and 147 new orders. While these numbers may be slightly inflated by incomplete genomes (see evaluation next section), it is clear that vConTACT3’s systematic approach rapidly and drastically impacts our understanding of current virus taxonomy. Supporting this, data from vConTACT3 has already been used to inform the selection and manual analysis of virus genomes in 18 family-rank taxonomic proposals submitted to the ICTV in 2024.

Finally, benchmarking against vConTACT2 and the only other scalable tools VPF-Class and geNomad revealed that vConTACT3 led or co-led in accuracy and other performance metrics, though all tools generally performed well (Supplemental Note 3). Again, while performance on “known” viruses was relatively similar, only vConTACT tools were able to create the new taxa needed to accommodate undocumented virus sequence space. As well, while the above is focused on prokaryotic viruses, performance benchmarking against ICTV was also done for eukaryotic viruses (13,524 from NCBI’s curated RefSeq, v220, accessed Nov. 2023), and found 96%, 95.9%, 96.7%, 98.7%, 76%, 84.4%, 100% concordance for genus, subfamily, family, order, class, phylum, realm, respectively (Extended Data Table 1). While we have not manually assessed these further to detail areas of discordance, the overwhelmingly high accuracy at most ranks suggests vConTACT3’s readiness for implementation for eukaryotic viruses, as well as targeted opportunities for ICTV taxonomic re-evaluations at class and phylum ranks.

### vConTACT3 learns from the known to taxonomically classify the unknown

As virus discovery studies now routinely observe thousands to hundreds of thousands of new viruses with most having no closely related reference genomes^45–52^, moving beyond reference-based classification with tools that can create new taxa is critical. Currently, non-vConTACT tools all rely on relationships against reference genomes. In contrast, vConTACT3 establishes rules by evaluating against ‘known’ sequence space using a statistical framework that allows it to extend these rules to unclassified sequence space. Thus, vConTACT3 can create new taxa to assign novel viruses – and does so at multiple taxonomic ranks. Given this unique, but important capability, we benchmarked it against the most significant challenge to taxonomy in virus discovery studies – genome fragmentation.

To examine this, we performed an *in silico* fragmentation experiment. Briefly, we randomly selected 20K IMG/VR virus sequences between 2K and 250K bp and fragmented them 10 times with fragment sizes Gaussian distributed and restricted to 20 - 80% of the original sequence’s length. This resulted in 41,536 total fragments, of which >90% (38,133) were classifiable by vConTACT3 at some rank (the remaining unclassifiable fragments did not have 3 or more genes shared with other genomes, **Fig. 4**). Larger fragments improved vConTACT3 placement, with most fragments (25,098, 65.8%) correctly placed through all hierarchical ranks down to genus, and the remaining also correctly placed, but under-classified such that ranks could only go down to subfamily (20.7%), class (8.5%), order (2.7%) or family (2.2%) (**Fig. 4a**). No fragment was incorrectly classified, rather it was taxonomic specificity that changed with fragment size with ~40% of the original genome size being a key threshold to get to genus or subfamily assignments. Examining by fragment lengths revealed that 1-3 Kb fragments are rarely classified to the genus rank, about one-third (35.1%) of 3-10 kb fragments classified to genus, and >10 kb fragments, which has been the community standard for previous versions of the vConTACT tools^31,32^, classified nearly all (96.3%) to genus or subfamily (**Fig. 4b**).

**Figure 3.**
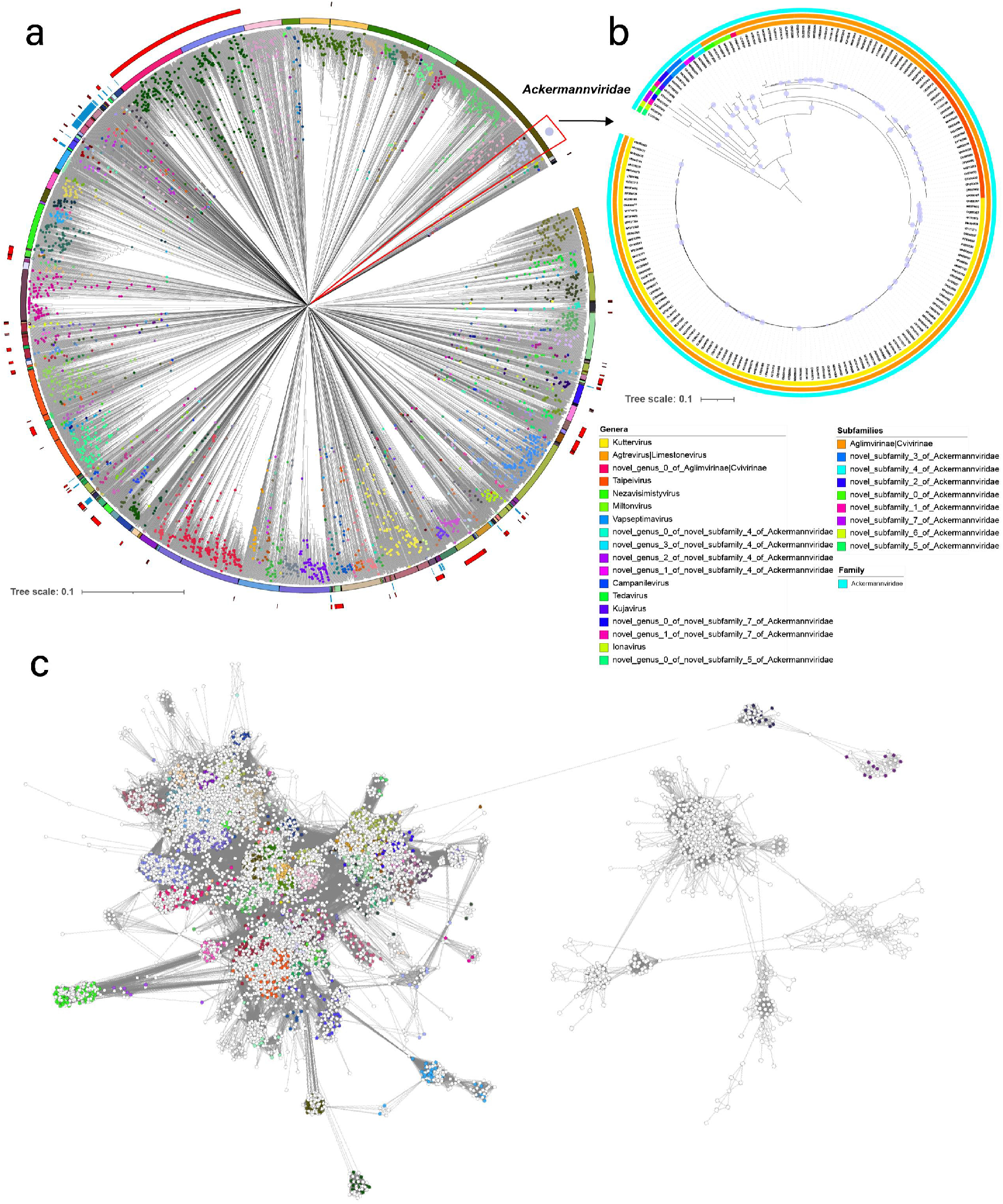
Comparing vConTACT3 versus ICTV taxonomic assignments. (**A**) Proteomic tree of 4,287 ICTV-classified prokaryotic and archaeal viruses annotated with vConTACT3-predicted orders and families. The tree was generated based on genomic distances from normalised tBLASTx scores using ViPTreeGen, visualised and annotated in iTOL. vConTACT3-predicted families are shown as coloured circles at branch midpoints, orders as coloured ranges in the first outer ring. Existing ICTV orders are depicted in blue in the second ring. Areas of discrepancy where vc3-predicted orders are split, or interspersed relative to the branch structure of the tree are denoted in red in the outermost ring. (**B**) Core-genome phylogeny of vConTACT3-predicted family corresponding to the *Ackermannviridae*. The tree was inferred with IQTree2 using trimmed MSAs of 8 core gene products. The tree is rooted at the midpoint and UFBoot branch support >95% is shown as filled circles. Genera, subfamilies and family predictions by vConTACT3 are demarcated as coloured outer rings. (**C**) Gene-sharing network of the 1Jun2024 INPHARED dataset as input comprising 23,227 sequences with nodes colored by order (corresponding to (**A**))

**Figure 4.**
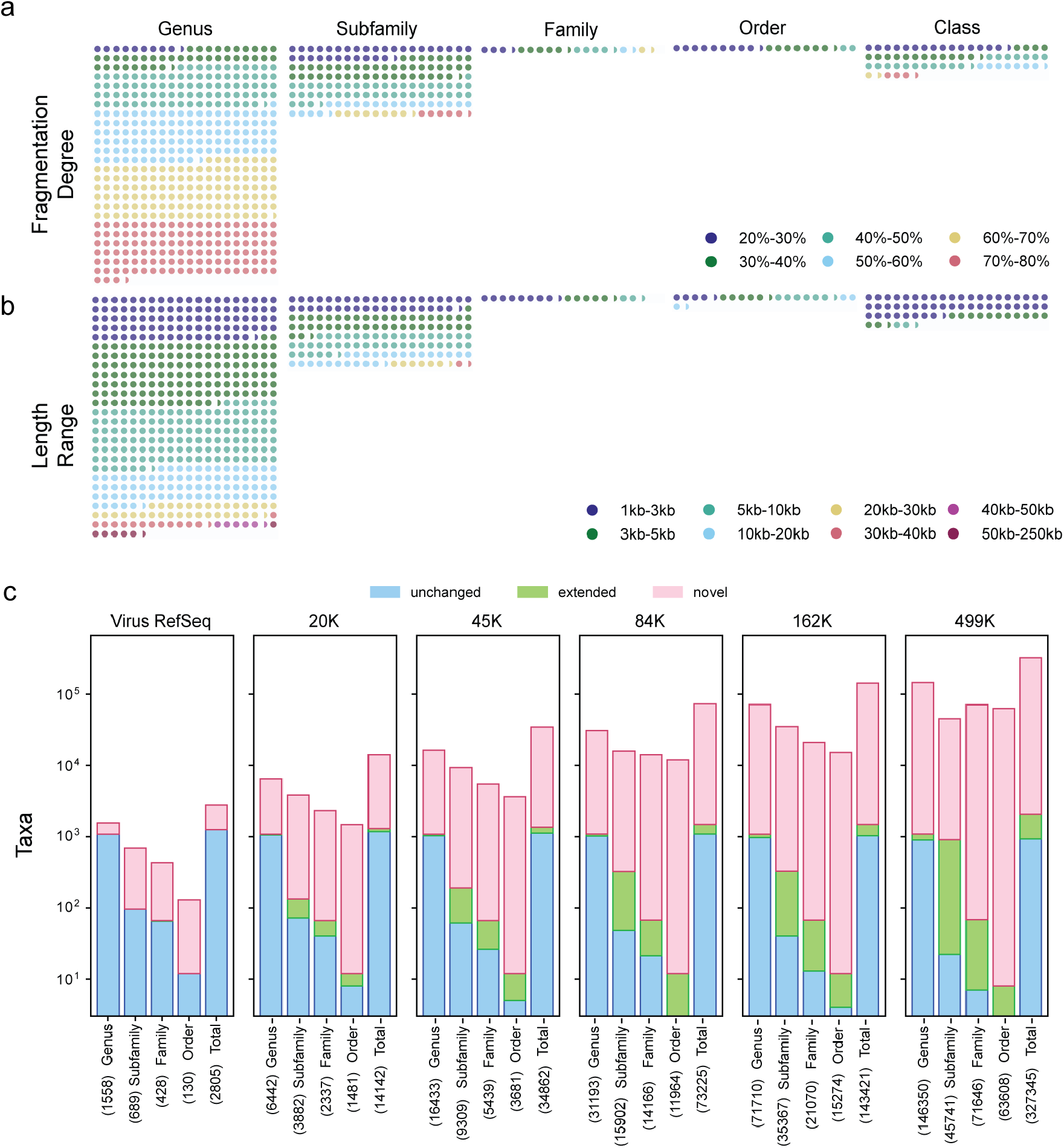
Assessment of the impact of genome fragments on taxonomic predictions. (a) Dot plot of IMG/VR fragments colored by the degree of fragmentation (20 - 80% of the original genomic fragment size), where each dot represents 50 fragments, and placement of the dots represents the best (i.e., highest) taxonomic rank that could be confidently assigned. (b) The same data as in A, but displayed by length instead of degree of fragmentation. (c) Comparison of vConTACT3 taxonomic assignment predictions against NCBI and with additions (20K sequences, 45K sequences, etc…) from IMG/VR. For each rank (genus, subfamily, family, and order), a taxon was considered *unchanged* if no genomes were added to its RefSeq representation, *extended* if new genomes were added, and *novel* if no reference taxonomy was available and a new taxon group needed to be created. The number of taxa represented is shown in parentheses.

Beyond fragment size, genomic content mattered for gene-sharing-based vConTACT3 assignments. For example, while the 3,654 genomic fragments that could not be assigned to any rank were predominantly less than 10 kb and thus might contain few genes (3,334 of 3,343), there were many examples where the right genomic content allowed correct assignments, as many 1-3 kb fragments classified fine when they contained key hallmark genes. For the 5,130 under-classified fragments (i.e., only to family, order, class instead of subfamily or genus), they varied between 3 and 59 genes (average = 5.3, median = 4.0) with these fragments sharing fewer genes with other genomes. Again these data suggest that the kinds of genes available on the genome fragment also matter. In summary, vConTACT3 correctly places partial genome fragments within their appropriate rank, but rank-level resolution depends upon fragment length and particular genomic context. While intuitive, the above now provides quantitative benchmarks for the interpretation of field data.

### Future outlook and conclusions

The ICTV, working closely with the global virology community, has leveraged a tiny window into Earth’s virosphere to create a taxonomic framework that strives to capture the evolutionary biology inherent to viruses – in spite of wildly different lifestyles and genomes that viruses can have. Through extensive benchmarking, we demonstrate that vConTACT3 accurately recapitulates ICTV taxonomy for known viruses, helps identify areas where patchwork taxonomic cut-offs might need revision, and provides a statistically grounded and systematic framework to extrapolate from known viruses into the vast regions of the virosphere that are currently unknown (**Fig. 4c**). In contrast to its predecessor, vConTACT3 also does so across many hierarchical ranks, across multiple realms, and at scales relevant to current large-scale sequencing surveys. Developed in tandem with ICTV members, vConTACT3 is proven here to be foundational for systematically formalizing taxonomic proposals, helping elucidate relationships in challenging areas of the virosphere, and automatically providing systematic multi-rank taxonomic assignments for tens of thousands of under-classified viruses already captured in public databases. Mindful that viruses across realms play by different evolutionary rules, vConTACT3 has been designed such that thresholds - currently implemented at the realm and host domain level - can be easily adjusted to incorporate experts’ feedback, either from updates to the taxonomy or from direct community feedback at the vConTACT3 Bitbucket repository. Together, these advances provide another large step forward in bringing big data to ICTV’s multi-rank vision^14^ and, we hope, more firmly establish genome-based approaches^16,18^ into the toolkit needed for viral taxonomists to handle classifying the vast and largely unexplored Earth’s virosphere.

## Acknowledgments

This work was supported by the National Science Foundation under Grants No. DBI-2149506 and DBI-2022070 (BII-Implementation: The EMERGE Institute) with high-performance computational support from the Ohio Supercomputer Center, the Deutsche Forschungsgemeinschaft (DFG, German Research Foundation) under Germany’s Excellence Strategy – EXC 2051 – Project-ID 390713860, the European Research Council (ERC) Consolidator grant 865694: DiversiPHI, the Alexander von Humboldt Foundation in the context of an Alexander von Humboldt-Professorship founded by German Federal Ministry of Education and Research, and the European Union’s Horizon 2020 research and innovation program, under the Marie Skłodowska-Curie Actions Innovative Training Networks grant agreement no. 955974 (VIROINF). E.M.A. gratefully acknowledges the support of the Biotechnology and Biological Sciences Research Council (BBSRC); this research was funded by the BBSRC Institute Strategic Programme Food Microbiome and Health BB/X011054/1 and its constituent projects BBS/E/F/000PR13631 and BBS/E/F/000PR13633; and by the BBSRC Institute Strategic Programme Microbes and Food Safety BB/X011011/1 and its constituent projects BBS/E/F/000PR13634, BBS/E/F/000PR13635 and BBS/E/F/000PR13636.

## Author Contributions

B.B. and M.B.S designed the study. B.B., O.Z., M.B.S wrote the manuscript with substantial contributions from all authors. D.T., H.B.J and B.B. performed the phylogenetic analyses and B.E.D., D.T., H.B.J, and B.B. performed the statistical and network analyses. J.G. and B.B. evaluated the distance metrics. B.B. developed the code with contributions from J.G.

## Code Availability

vConTACT3 is available through Bitbucket (https://bitbucket.org/MAVERICLab/vcontact3) as an installable Python package, as well as through python package managers Anaconda (https://anaconda.org/) and Mamba (https://mamba.readthedocs.io/). Instructions for building an Apptainer container of vConTACT3 is available on Bitbucket, along with a definitions file.

Databases used by vConTACT3 are available as citable datasets at Zenodo (https://zenodo.org/) under https://doi.org/10.5281/zenodo.10035619 and https://doi.org/10.5281/zenodo.10935513.

## Methods

### Methodology

vConTACT3 takes a set of genomes, predicts their gene sequences with pyrodigal^53^ (or other options), passes these genes to MMSeqs2, and clusters them at 30% - 70% minimum AAI in 10% increments. This creates protein clusters (PC), which represent all of the genes that are similar under that identity. Independently, each of these 5 clustering identities are used to build a PC profile via SciPy^54^‘s crosstab function, which results in a genome X PC matrix. Each PC profile matrix is then used to construct a distance matrix between genomes using a vectorized version of the “SqRoot” distance metric (see Methods). This distance matrix is converted into a network using the networkit^55^ package, and filtered to remove edges with fewer than 3 genes shared. References within each network component are first used to assign realm-level taxonomy. In cases where two or more realms are identities within the same component, the minimum shared genes within the component are increased until the multi-realm membership of the component are severed. Removed edges are added back in, unless they would otherwise re-connect different realms. For some realms (e.g. *Adnaviria* and *Varidnaviria*), separation is deemed impossible, so “hybrid” realm distances are used and applied to the entire component. After this multi-realm removal step, networks are converted into a matrix and agglomerative clustering performed on each of the network’s connected components using SciPy’s hierarchy function, using the Euclidean metric and average method for new clusters. This linkage matrix is then separated into flat clusters using pairwise distance cutoffs determined for each virus realm and host domain by benchmarking (see Parameter Optimization). This results in a set of assignments for each genome, which are aggregated as a result of their optimal clustering identity and pairwise cutoff. For example, genomeA may be in a cluster deriving from 30% identity at the order level, a cluster from 50% at family level, and two clusters from 70% at subfamily and genus levels (differing in pairwise distance cutoffs). Each cluster is assigned a classification depending on its rank, leveraging reference genomes if they co-occur in the same cluster. This assignment is performed independently at each rank, and allows situations where a 7-member order can have an 8-member family directly underneath, reflecting the inconsistencies within the ICTV taxonomy’s distances (e.g. where average distances between family A may be markedly different than distances between family B within the same realm). In situations like these, assignments are given “||” indicators, allowing users to quickly identify these regions. Additionally, where reference taxonomies exist, it is possible for multiple references of differing classifications to exist within the same cluster, in which case genomes have their taxons merged and assigned “|”. New taxonomy assignments (where no references exist) are created using the “novel_<rank>_<#>_of_<upper-rank>” template, where <rank> signifies the rank of the taxon being created, <#> the count of newly created ranks, and <upper-rank> signifies the rank at the higher taxonomic level.

### Reference datasets for benchmarking

To construct the benchmarking dataset and reference database, proteins from NCBI Virus RefSeq version 218 were downloaded, along with the RefSeq Release 218 catalog (consisting of protein to genome accession mapping), as well as the virus genome reports file (containing NCBI genome info). Additionally, the latest version of the VirusHostDB was downloaded. The VirusHostDB and the virus genome reports file were merged on taxon id. Viruses with segmented or multipartite genomes were combined in a single record, viruses with strain information had their representative genome selected (discarding others), and viruses without host information were excluded. Viruses were then annotated with full taxonomy using the ETE3 ToolKit^56^, and missing realm information was filled in using the NCBI virus lineage field from the genome report. All ranks were kept, except for subphylum, suborder, and subgenus. In cases where no information was provided, Entrez queries were constructed to fill in, wherever possible. This resulted in a table containing 21,080 genomes spanning all 6 virus realms and all 3 host domains. Finally, virus protein sequences were filtered against this table to remove those from viruses without host or virus realm information, or from discarded strains. These final proteins were then built into separate datasets according to their host domain, excluding Archaea (due to so few genomes) and an additional “prokaryotes” dataset, merging both Archaea and Bacteria hosts.

### Parameter optimization

In order to exhaustively benchmark vConTACT3, each of the 3 major steps (PC profile generation, gene-sharing metric, and hierarchical clustering) was permuted for all 3 domain datasets (*Eukaryota* and prokaryotes). For PC profile generation, this included clustering using MMSeqs2^57^ version 14-7e284 from 30 - 70% in 10% increments, and 80, 90, and 95% clustering identities using the “--min-seq-id” parameter, keeping all other parameters as default. For the gene-sharing metric, used to define the distance between genomes, 4 metrics were selected; 1) the Jaccard, 2) the “VirClust” metric, used by VirClust 3) “Shorter”, and 4) “SqRoot”, shown below:

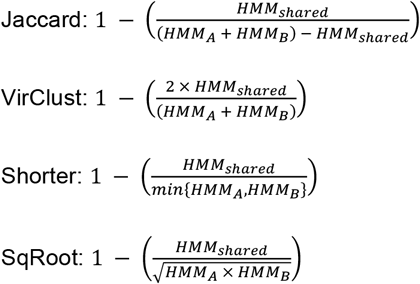

where *HMM*_*shared*_ are the number of PCs shared between genomes A and B, *HMM*_*A*_ are the number of PCs in genome A, and *HMM*_*B*_ are the number of PCs in genome B. Minimum PCs required to be shared between genomes were also assessed (known as “shared genes”), ranging from 1, 2, 3, 4, 5, 10, 15, 20, and 25. For hierarchical clustering, following agglomerative clustering using SciPy’s linkage function using the “average” method, clusters were determined by “fcluster”, also part of the SciPy package, using criterion of “distance”, and ranged between the values of 0 and 1 in 0.01 increments.

In total, 8 clustering identities were used to assess PC profile generations, 4 metrics to define genome distances, 8 minimum shared gene cutoffs, and 100 hierarchical clustering distance cutoffs against 3 host domains, and for all 6 virus realms. Additionally, since the networks used to define the span of hierarchical clusters are separated by connected components, *each* component was also evaluated. This resulted in 38,117,688 and 22,425,924 combinations for prokaryotes and eukaryotes, respectively, and excluded connected components with 2 or fewer genomes. Th*/e resulting taxonomic assignments were evaluated using traditional performance metrics, such as accuracy, separation, sensitivity and positive predictive value (PPV), as well as network and clustering-based approaches, such as the adjusted Rand index, and mutual information. Central to accurate classification was rigorously evaluating each step of the updated methodology against the NCBI reference taxonomy. An intentional choice was chosen regarding the use of NCBI’s taxonomy over ICTV, as the ICTV taxonomy does not cover the breadth of sequences in NCBI, and merging both taxonomies introduces situations where parts of taxon ranks have been updated/evaluated by ICTV whereas others are not, leading to different assignments.

In order to select the best combination of parameters, the >60 million combinations were grouped separately by host domain (3) and virus realm (6), grouped by rank within each realm (realm selections were handled independently), and then sorted by accuracy and ARI. Instead of automatically selecting the maximum values, accuracy plots (Extended Data Fig. 2-7 and Fig. 2) were used to guide by allowing visual inspection of the curve slopes. Most of the lower ranks (genus, subfamily, and family) had slopes with clearly defined maxima or plateau. For the latter, multiple optimal solutions were available, and were prioritized as follows: 1) selecting clustering identities with a greater proportion of highest accuracies and ARIs relative to other identities, 2) selecting pairwise distances that maximized the distance between selection of other ranks (i.e. genus and subfamily both using 70% identity and genus plateaus between 0.50 - 0.70 and subfamily between 0.60 - 0.75, subfamily would be >0.7 and genus would be <0.6), and 3) falling back to sensitivity, PPV, and other measures. In cases where all other factors were equal, the mean of the cutoffs were selected. Given the semi-supervised selection of parameters, it is possible for more optimal solutions to exist, but were not pursued due to many selections providing >=97% accuracy. Of particular note - despite independent selection - several virus realms “matched” each other under the exact same set of parameters. For the upper ranks (class, phylum, kingdom), accuracies were greatly reduced and generally trended with higher accuracy as cutoffs approached 1. Additionally, while all were evaluated, only shared genes of 3 or more were selected as that was the threshold determined (see below) to eliminate cross-realm edges.

### Performance comparisons

Scalability benchmarks between v3, geNomad and VPF-Class were conducted using a subset (n=5,400,001) of IMG/VR v4.1^15^ (December 2022 release) virus sequences, and measured at multiple sample fractions (0.1%, 0.25%, 0.5%, 1.0%, 2.5%, 5%, 10%, 15%, 20% - 100% (10% increments). Sub-1% fractions were done to allow vConTACT2 to be sufficiently benchmarked before its software limitations were encountered, and for other tools to show upper limits regarding scale. The final set of virus sequences used was derived by first obtaining the ‘full’ set (n = 15,677,623), and then filtering to exclude genomes under 2,000 bp (n = 15,505,302), greater than 250 kb (n = 15,500,802), and keeping only “high-confidence” assignments, totaling 5,400,001 virus sequences. We note that although we wanted to include VirClust in our performance comparisons, we were unable to do so since its design and code meant it would have taken significant time and resources to bring it to comparable scale to vConTACT, VPF-Class and geNomad.

### Fragmentation analysis

From the 0.1% scalability dataset, sequences were randomly fragmented using a custom python script as follows: a genome fraction percentage (i.e. 27%, 35%, 55%, etc) was randomly selected using the numpy package’s random choice using a uniform distribution. This was converted to a unit of genome length (27% of 10 kb = 2.7 kb) and then passed to numpy’s randomized choice using a normal (e.g. Gaussian) distribution as the mean selection value with a standard deviation of 10%, resulting in 10 total lengths (). The start positions of these 10 lengths along the genome were randomly selected using random int (start position from 0 to genome length). In cases where the start position would be placed at a point where start position + genome length would exceed the total genome length (start position of 9000 with a 2.7 kb length > 10 kb full genome), the fragment was discarded. Additionally, fragment lengths were required to be greater than 20% of the genome’s original length as well as less than 80%, assuming that 400-bp would be sufficient for a single gene (20% of a 2 kb genome) and 80% representing the majority-not-full-length of a genome. Though a single gene was assumed, no test was performed to ensure a minimum number of potential genes. The result of the fragmentation process yielded 41,536 total fragments.

To assess the classification specificity, fragmented genomes were processed twice with vConTACT3, once including the “reference” original genome alongside their fragments, and once excluding the original genome. The original-excluding dataset was performed to reduce bias towards fragmented genomes aligning with their full-length counterpart. The original-including genome groups of each rank (e.g. class, order, family, subfamily, genus) were compared against the original-excluding ones (genusA_of_full_length vs genusZ_of_fragmented). When differences occurred between members of the same group (relative to the references), the rank of most agreement was selected.

### ML-Phylogeny

To assess inter- and intra-PCs similarity at genus, subfamily and family levels, a total of 10 families were selected from three vc3-predicted orders. For each family, PCs were identified using MMSeqs2^57^ at 30% minimum identity and 50% coverage. To assess phylogenetic support, partitioned maximum likelihood trees were calculated using PCs shared by >99% of genomes within the vc3-predicted family using IQTree2^58–60^ with 1000 ultra-fast bootstrap and SH-Alrt replicates, with -m TEST for selection of the best-fit model. Trees were mid-point rooted, visualized and annotated using iToL^61^. Intergenomic nucleotide similarities were calculated using taxmyPhage^62^. Annotated heatmaps of shared PCs and nucleotide similarities were produced using custom python scripts.

