## Extended Data Figures for "Scalable and systematic hierarchical virus taxonomy with vConTACT3"

**Running Title**: Multi-rank, large-scale virus classification with vConTACT3

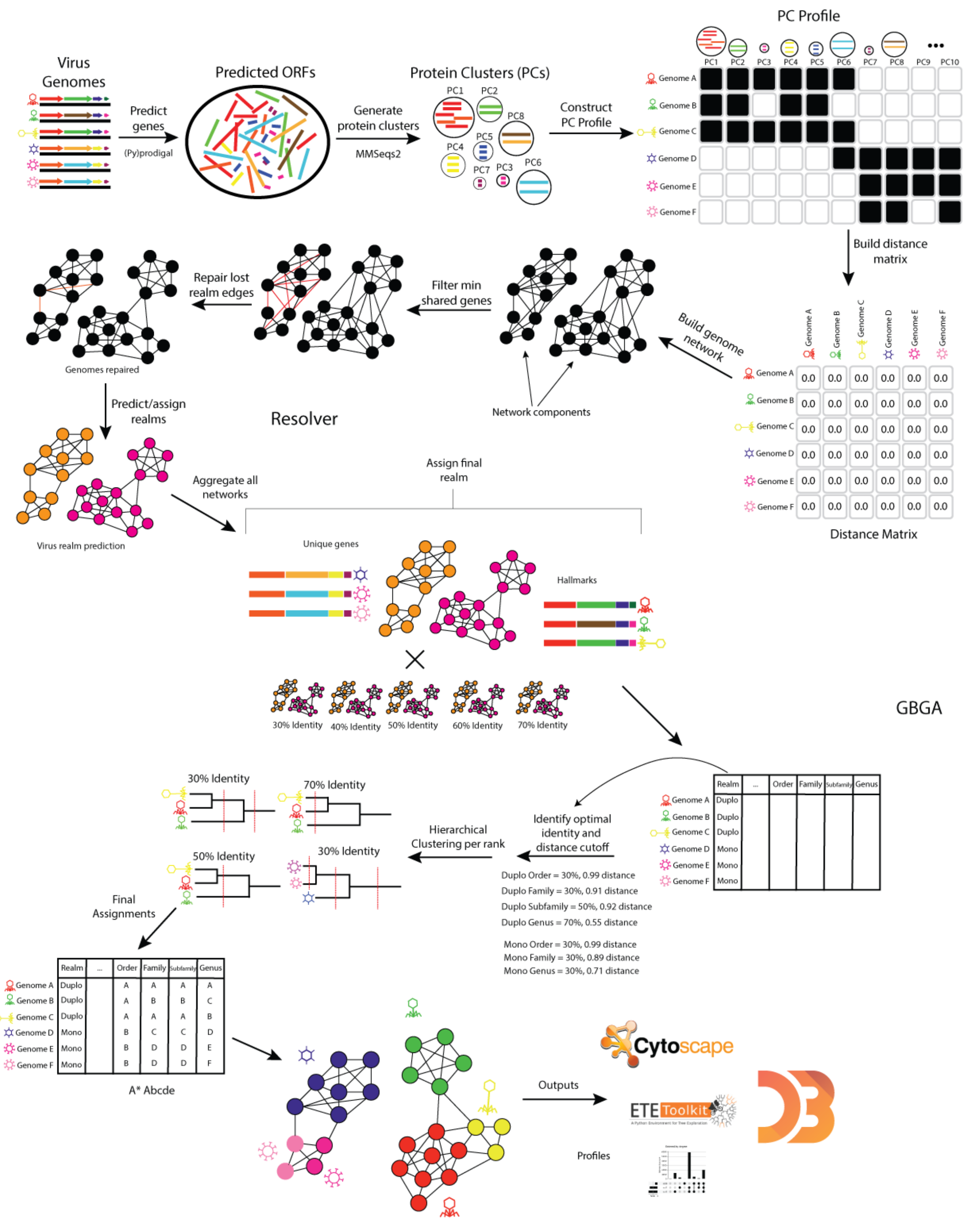


**Extended Data Fig. 1. Detailed vConTACT workflow and outputs.** User genomes are provided to vConTACT3, genes predicted with prodigal, and then sent to MMSeqs2 to be clustered at identities 30% - 70% in 10% increments. These 5 clustering identities are used to build a protein cluster (PC) profile, corresponding to each clustering identity. These 5 PC profiles are sent to the “resolver,” which constructs a distance matrix based on the selected distance metric (default “SqRoot”, see **Methods**). This distance matrix is subsequently converted into a network, which is annotated with any available genome information. The network is then filtered for the minimum number of shared genes allowed between genomes, and then “repaired” if they are within the same connected component. Additionally, users can select a high-accuracy repair, which more carefully reviews dropped edges. The final stage of the resolver predicts virus realms the genomes belong to, using network edge-connected references and/or the presence of PCs exclusively identified and co-shared with references. Entirely novel genomes are assigned a default, user-selectable realm. The 5 filtered and repaired networks are then sent to the guilt-by-genome-association (GBGA) assiger. The GBGA aggregates realm predictions per-genome and per-network component and assigns a final realm for each genome. This realm prediction is used to select the optimal distance cutoff for each virus rank (genus, subfamily, family and order) and hierarchically cluster all genomes of that realm at that cutoff. These clusters are then matched against references (if co-clustered and/or available) or used as novel ranks to assign each virus rank order and below. (The upper ranks of phylum and class inherit reference-based assignments within the predicted realm). After assignments, GBGA output is then compared with reference sequences (if available) and performance metrics are calculated. Finally, PC profiles, performance metrics, and GBGA outputs are integrated into the exports/results component, which provides user-controlled outputs in Cytoscape format, a d3js interactive HTML network, UpSet plots, profiles, and Newick-formatted dendrograms.

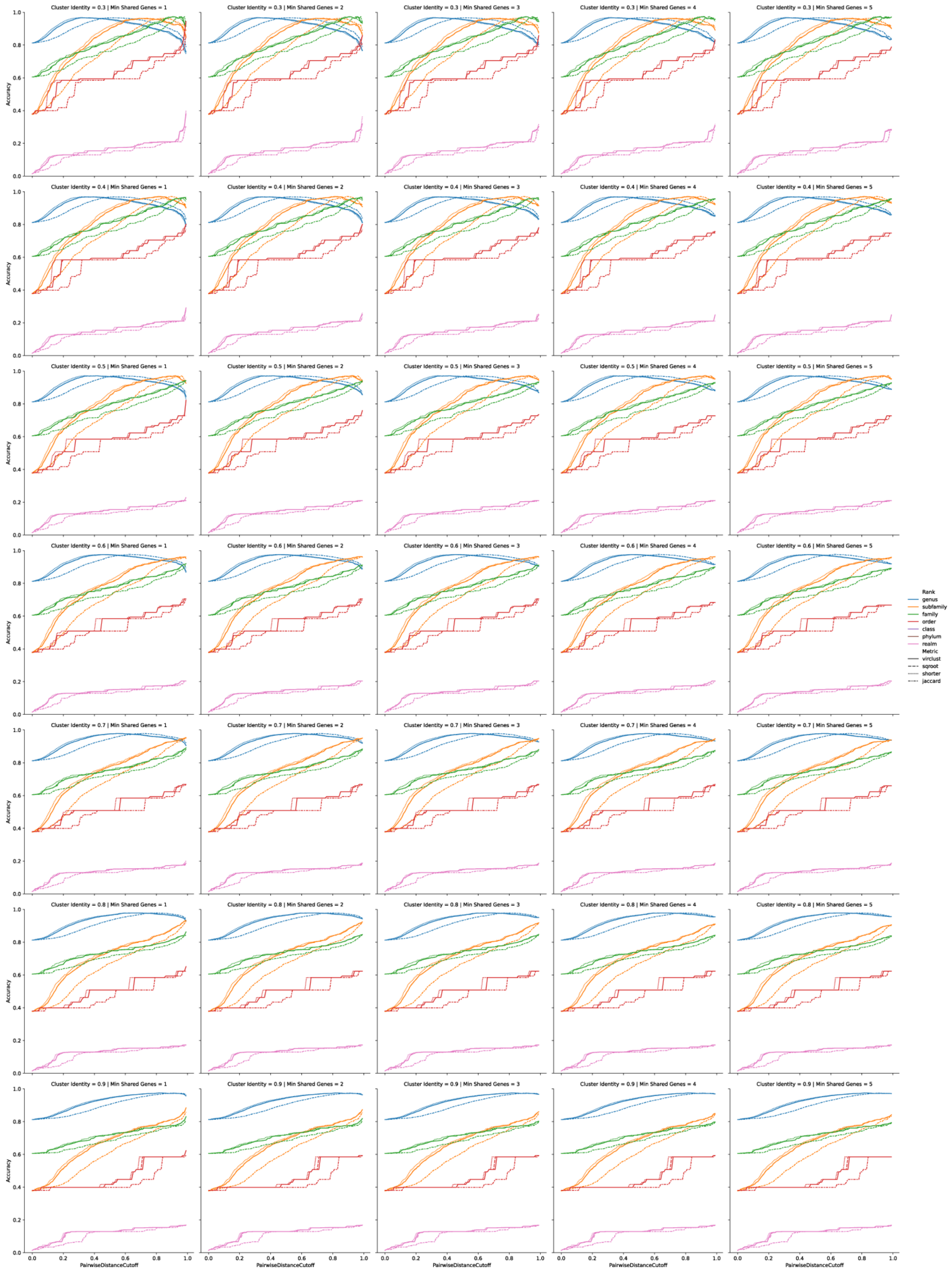
Cluster Identity = 0.3 | Min Shared Genes = 1
Cluster Identity = 0.3 | Min Shared Genes = 2
Cluster Identity = 0.3 | Min Shared Genes = 3
Cluster Identity = 0.3 | Min Shared Genes = 4
Cluster Identity = 0.3 | Min Shared Genes = 5
Cluster Identity = 0.4 | Min Shared Genes = 1
Cluster Identity = 0.4 | Min Shared Genes = 2
Cluster Identity = 0.4 | Min Shared Genes = 3
Cluster Identity = 0.4 | Min Shared Genes = 4
Cluster Identity = 0.4 | Min Shared Genes = 5
Cluster Identity = 0.5 | Min Shared Genes = 1
Cluster Identity = 0.5 | Min Shared Genes = 2
Cluster Identity = 0.5 | Min Shared Genes = 3
Cluster Identity = 0.5 | Min Shared Genes = 4
Cluster Identity = 0.5 | Min Shared Genes = 5
Cluster Identity = 0.6 | Min Shared Genes = 1
Cluster Identity = 0.6 | Min Shared Genes = 2
Cluster Identity = 0.6 | Min Shared Genes = 3
Cluster Identity = 0.6 | Min Shared Genes = 4
Cluster Identity = 0.6 | Min Shared Genes = 5
Cluster Identity = 0.7 | Min Shared Genes = 1
Cluster Identity = 0.7 | Min Shared Genes = 2
Cluster Identity = 0.7 | Min Shared Genes = 3
Cluster Identity = 0.7 | Min Shared Genes = 4
Cluster Identity = 0.7 | Min Shared Genes = 5
Cluster Identity = 0.8 | Min Shared Genes = 1
Cluster Identity = 0.8 | Min Shared Genes = 2
Cluster Identity = 0.8 | Min Shared Genes = 3
Cluster Identity = 0.8 | Min Shared Genes = 4
Cluster Identity = 0.8 | Min Shared Genes = 5
Cluster Identity = 0.9 | Min Shared Genes = 1
Cluster Identity = 0.9 | Min Shared Genes = 2
Cluster Identity = 0.9 | Min Shared Genes = 3
Cluster Identity = 0.9 | Min Shared Genes = 4
Cluster Identity = 0.9 | Min Shared Genes = 5
Accuracy
PairwiseDistanceCutoff
Rank
genus
subfamily
family
order
class
phylum
realm
Metric
virclust
sqroot
shorter
jaccard

**Extended Data Fig. 2. Prokaryotic-infecting viruses of the *Duplornaviria* realm.** A 5x7 grid of line plots representing accuracies of *Duplornaviria* viruses infecting prokaryotes (*Bacteria* and *Archaea*). From left-to-right, plots increase in the minimum number of shared genes required (ranging from 1 - 5) between genomes to be considered as related. From top-to-bottom, plots increase in minimum clustering identity used to establish protein clusters (PCs). Since PCs are used to determine the number of shared genes between genomes, increasing clustering identity increases the stringency required for two genes between two separate genomes to be considered shared, and thus, related. Within each plot, accuracy (Y-axis) is a measure of agreement between NCBI taxonomy and vConTACT3 predictions. Pairwise distance cutoff (X-axis) represents the cutoff threshold used during hierarchical clustering to define clusters. The cutoff ranges between 0 - 0.99, with 0 representing completely identical PC profiles, and 0.99 representing nearly no shared genes. Line colors represent taxonomic rank, and dashed lines represent the type of distance metric (Jaccard, “SqRoot”, ‘VirClust” and “Shorter”) employed. See **Methods** for description of each distance metric.

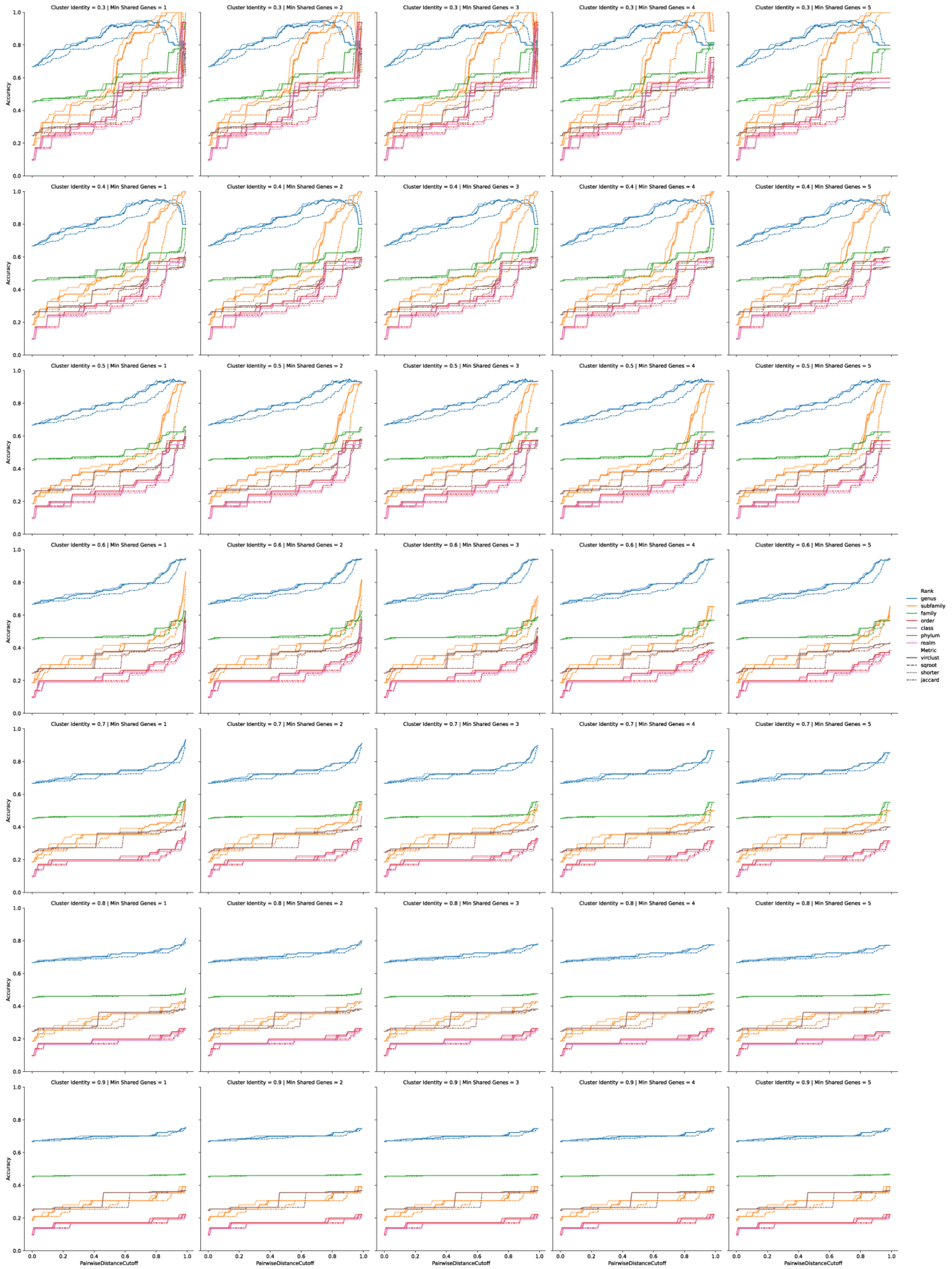
Cluster Identity = 0.3 | Min Shared Genes = 1
Cluster Identity = 0.3 | Min Shared Genes = 2
Cluster Identity = 0.3 | Min Shared Genes = 3
Cluster Identity = 0.3 | Min Shared Genes = 4
Cluster Identity = 0.3 | Min Shared Genes = 5
Cluster Identity = 0.4 | Min Shared Genes = 1
Cluster Identity = 0.4 | Min Shared Genes = 2
Cluster Identity = 0.4 | Min Shared Genes = 3
Cluster Identity = 0.4 | Min Shared Genes = 4
Cluster Identity = 0.4 | Min Shared Genes = 5
Cluster Identity = 0.5 | Min Shared Genes = 1
Cluster Identity = 0.5 | Min Shared Genes = 2
Cluster Identity = 0.5 | Min Shared Genes = 3
Cluster Identity = 0.5 | Min Shared Genes = 4
Cluster Identity = 0.5 | Min Shared Genes = 5
Cluster Identity = 0.6 | Min Shared Genes = 1
Cluster Identity = 0.6 | Min Shared Genes = 2
Cluster Identity = 0.6 | Min Shared Genes = 3
Cluster Identity = 0.6 | Min Shared Genes = 4
Cluster Identity = 0.6 | Min Shared Genes = 5
Cluster Identity = 0.7 | Min Shared Genes = 1
Cluster Identity = 0.7 | Min Shared Genes = 2
Cluster Identity = 0.7 | Min Shared Genes = 3
Cluster Identity = 0.7 | Min Shared Genes = 4
Cluster Identity = 0.7 | Min Shared Genes = 5
Cluster Identity = 0.8 | Min Shared Genes = 1
Cluster Identity = 0.8 | Min Shared Genes = 2
Cluster Identity = 0.8 | Min Shared Genes = 3
Cluster Identity = 0.8 | Min Shared Genes = 4
Cluster Identity = 0.8 | Min Shared Genes = 5
Cluster Identity = 0.9 | Min Shared Genes = 1
Cluster Identity = 0.9 | Min Shared Genes = 2
Cluster Identity = 0.9 | Min Shared Genes = 3
Cluster Identity = 0.9 | Min Shared Genes = 4
Cluster Identity = 0.9 | Min Shared Genes = 5
Accuracy
PairwiseDistanceCutoff
Rank
genus
subfamily
family
order
class
phylum
realm
Metric
virclust
sqroot
shorter
jaccard

**Extended Data. Fig 3. Eukaryotic-infecting viruses of the *Duplornaviria* realm.** A 5x7 grid of line plots representing accuracies of *Duplornaviria* viruses infecting Eukaryota. Details are as Extended Data Fig. 2.

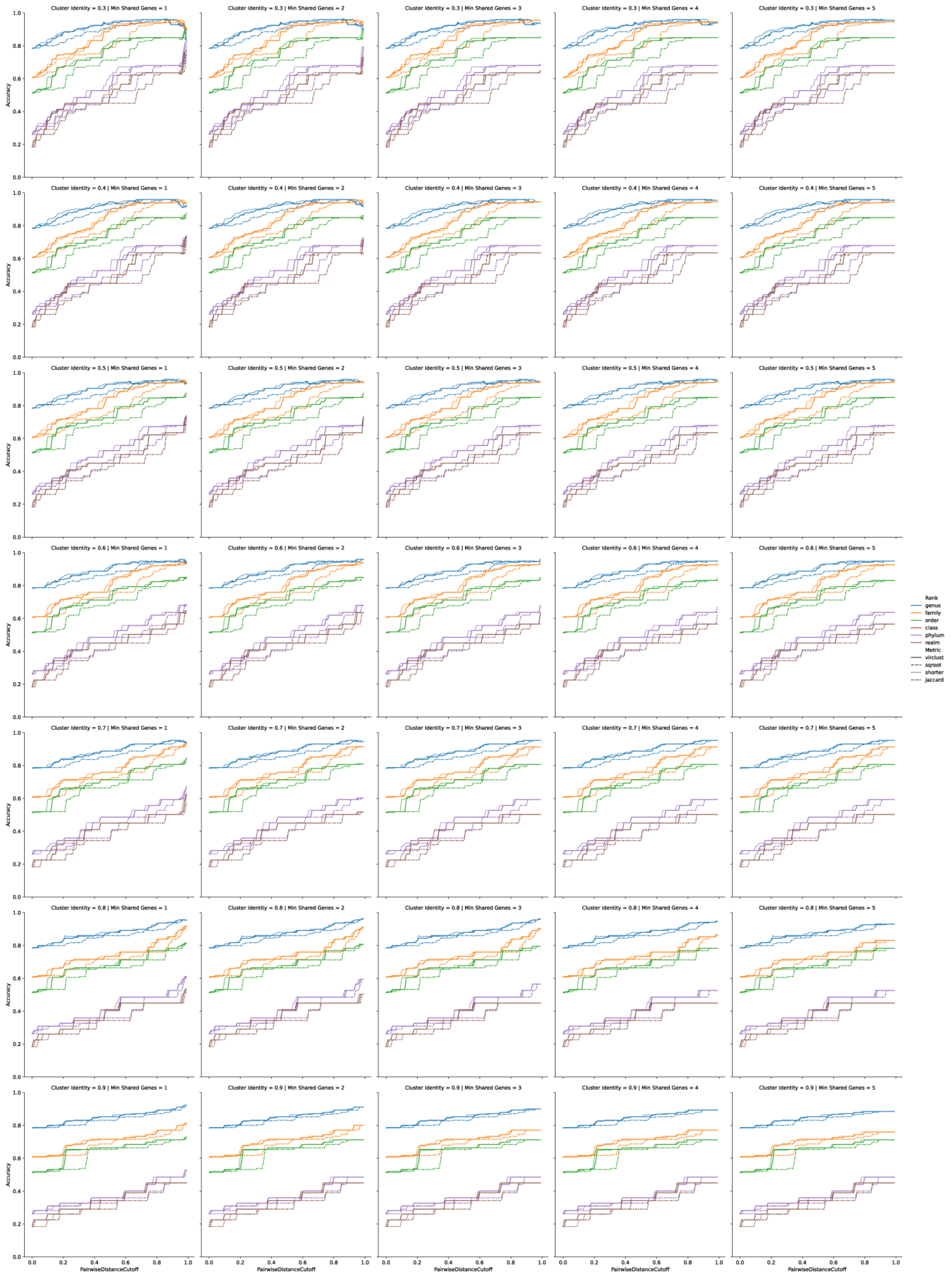

Cluster Identity = 0.3 | Min Shared Genes = 1
Cluster Identity = 0.3 | Min Shared Genes = 2
Cluster Identity = 0.3 | Min Shared Genes = 3
Cluster Identity = 0.3 | Min Shared Genes = 4
Cluster Identity = 0.3 | Min Shared Genes = 5
Cluster Identity = 0.4 | Min Shared Genes = 1
Cluster Identity = 0.4 | Min Shared Genes = 2
Cluster Identity = 0.4 | Min Shared Genes = 3
Cluster Identity = 0.4 | Min Shared Genes = 4
Cluster Identity = 0.4 | Min Shared Genes = 5
Cluster Identity = 0.5 | Min Shared Genes = 1
Cluster Identity = 0.5 | Min Shared Genes = 2
Cluster Identity = 0.5 | Min Shared Genes = 3
Cluster Identity = 0.5 | Min Shared Genes = 4
Cluster Identity = 0.5 | Min Shared Genes = 5
Cluster Identity = 0.6 | Min Shared Genes = 1
Cluster Identity = 0.6 | Min Shared Genes = 2
Cluster Identity = 0.6 | Min Shared Genes = 3
Cluster Identity = 0.6 | Min Shared Genes = 4
Cluster Identity = 0.6 | Min Shared Genes = 5
Cluster Identity = 0.7 | Min Shared Genes = 1
Cluster Identity = 0.7 | Min Shared Genes = 2
Cluster Identity = 0.7 | Min Shared Genes = 3
Cluster Identity = 0.7 | Min Shared Genes = 4
Cluster Identity = 0.7 | Min Shared Genes = 5
Cluster Identity = 0.8 | Min Shared Genes = 1
Cluster Identity = 0.8 | Min Shared Genes = 2
Cluster Identity = 0.8 | Min Shared Genes = 3
Cluster Identity = 0.8 | Min Shared Genes = 4
Cluster Identity = 0.8 | Min Shared Genes = 5
Cluster Identity = 0.9 | Min Shared Genes = 1
Cluster Identity = 0.9 | Min Shared Genes = 2
Cluster Identity = 0.9 | Min Shared Genes = 3
Cluster Identity = 0.9 | Min Shared Genes = 4
Cluster Identity = 0.9 | Min Shared Genes = 5
Accuracy
0.0
0.2
0.4
0.6
0.8
1.0
PairwiseDistanceCutoff
Rank
genus
family
order
class
phylum
realm
Metric
virclust
sqroot
shorter
jaccard

**Extended Data Fig. 4. Prokaryotic-infecting viruses of the Adnaviria and Varidnaviria realms**. A 5x7 grid of line plots representing accuracies of *Adnaviria and Varidnaviria* viruses infecting prokaryotes (*Bacteria* and *Archaea*). Details are as Extended Data Fig. 2.

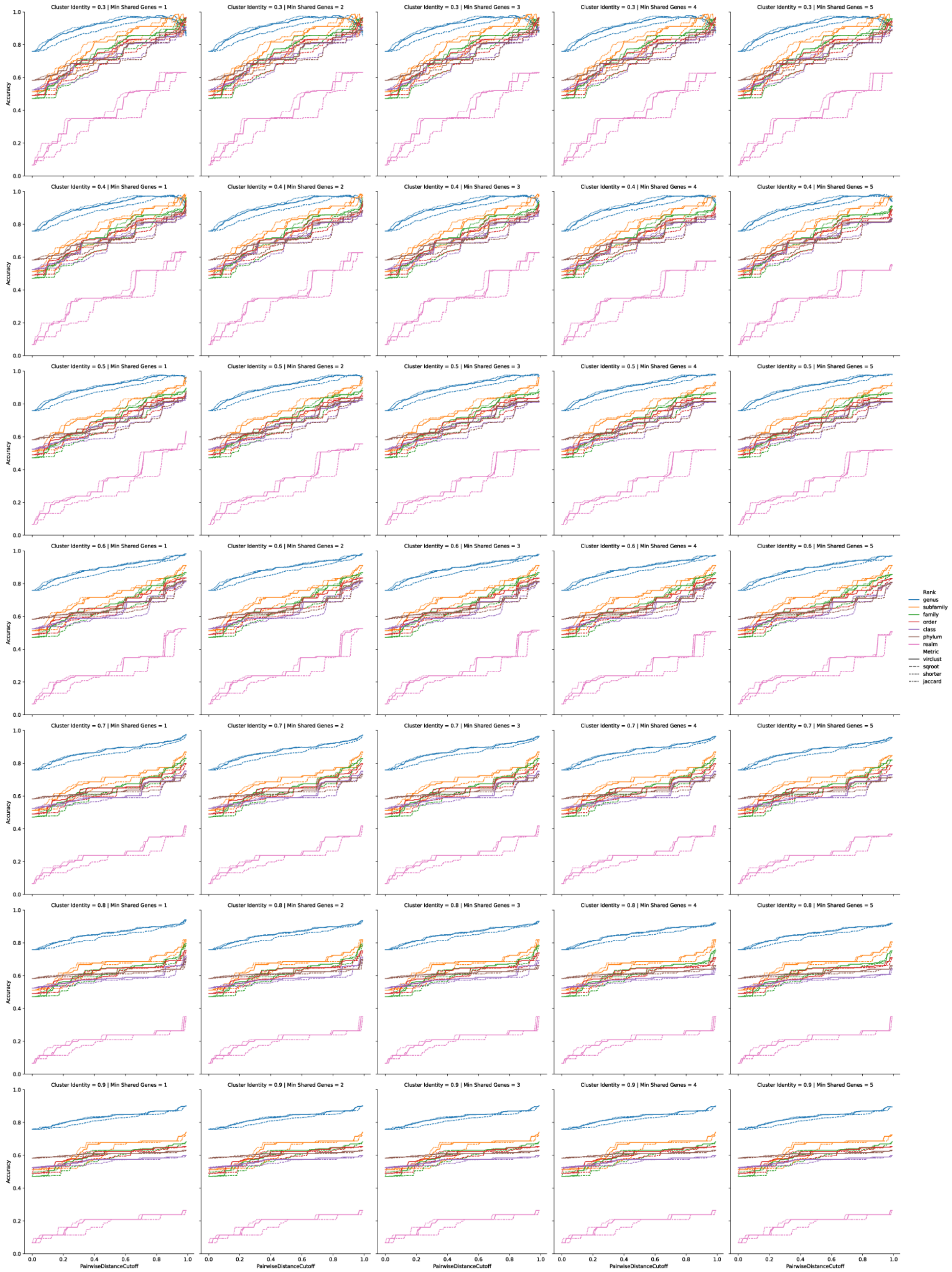
Cluster Identity = 0.3 | Min Shared Genes = 1
Cluster Identity = 0.3 | Min Shared Genes = 2
Cluster Identity = 0.3 | Min Shared Genes = 3
Cluster Identity = 0.3 | Min Shared Genes = 4
Cluster Identity = 0.3 | Min Shared Genes = 5
Cluster Identity = 0.4 | Min Shared Genes = 1
Cluster Identity = 0.4 | Min Shared Genes = 2
Cluster Identity = 0.4 | Min Shared Genes = 3
Cluster Identity = 0.4 | Min Shared Genes = 4
Cluster Identity = 0.4 | Min Shared Genes = 5
Cluster Identity = 0.5 | Min Shared Genes = 1
Cluster Identity = 0.5 | Min Shared Genes = 2
Cluster Identity = 0.5 | Min Shared Genes = 3
Cluster Identity = 0.5 | Min Shared Genes = 4
Cluster Identity = 0.5 | Min Shared Genes = 5
Cluster Identity = 0.6 | Min Shared Genes = 1
Cluster Identity = 0.6 | Min Shared Genes = 2
Cluster Identity = 0.6 | Min Shared Genes = 3
Cluster Identity = 0.6 | Min Shared Genes = 4
Cluster Identity = 0.6 | Min Shared Genes = 5
Cluster Identity = 0.7 | Min Shared Genes = 1
Cluster Identity = 0.7 | Min Shared Genes = 2
Cluster Identity = 0.7 | Min Shared Genes = 3
Cluster Identity = 0.7 | Min Shared Genes = 4
Cluster Identity = 0.7 | Min Shared Genes = 5
Cluster Identity = 0.8 | Min Shared Genes = 1
Cluster Identity = 0.8 | Min Shared Genes = 2
Cluster Identity = 0.8 | Min Shared Genes = 3
Cluster Identity = 0.8 | Min Shared Genes = 4
Cluster Identity = 0.8 | Min Shared Genes = 5
Cluster Identity = 0.9 | Min Shared Genes = 1
Cluster Identity = 0.9 | Min Shared Genes = 2
Cluster Identity = 0.9 | Min Shared Genes = 3
Cluster Identity = 0.9 | Min Shared Genes = 4
Cluster Identity = 0.9 | Min Shared Genes = 5
Accuracy
PairwiseDistanceCutoff
Rank
genus
subfamily
family
order
class
phylum
realm
Metric
virclust
sqroot
shorter
jaccard

**Extended Data Fig. 5. Eukaryotic-infecting viruses of the Adnaviria and Varidnaviria realms**. A 5x7 grid of line plots representing accuracies of *Adnaviria and Varidnaviria* viruses infecting *Eukaryota*. Details are as Extended Data Fig. 2.

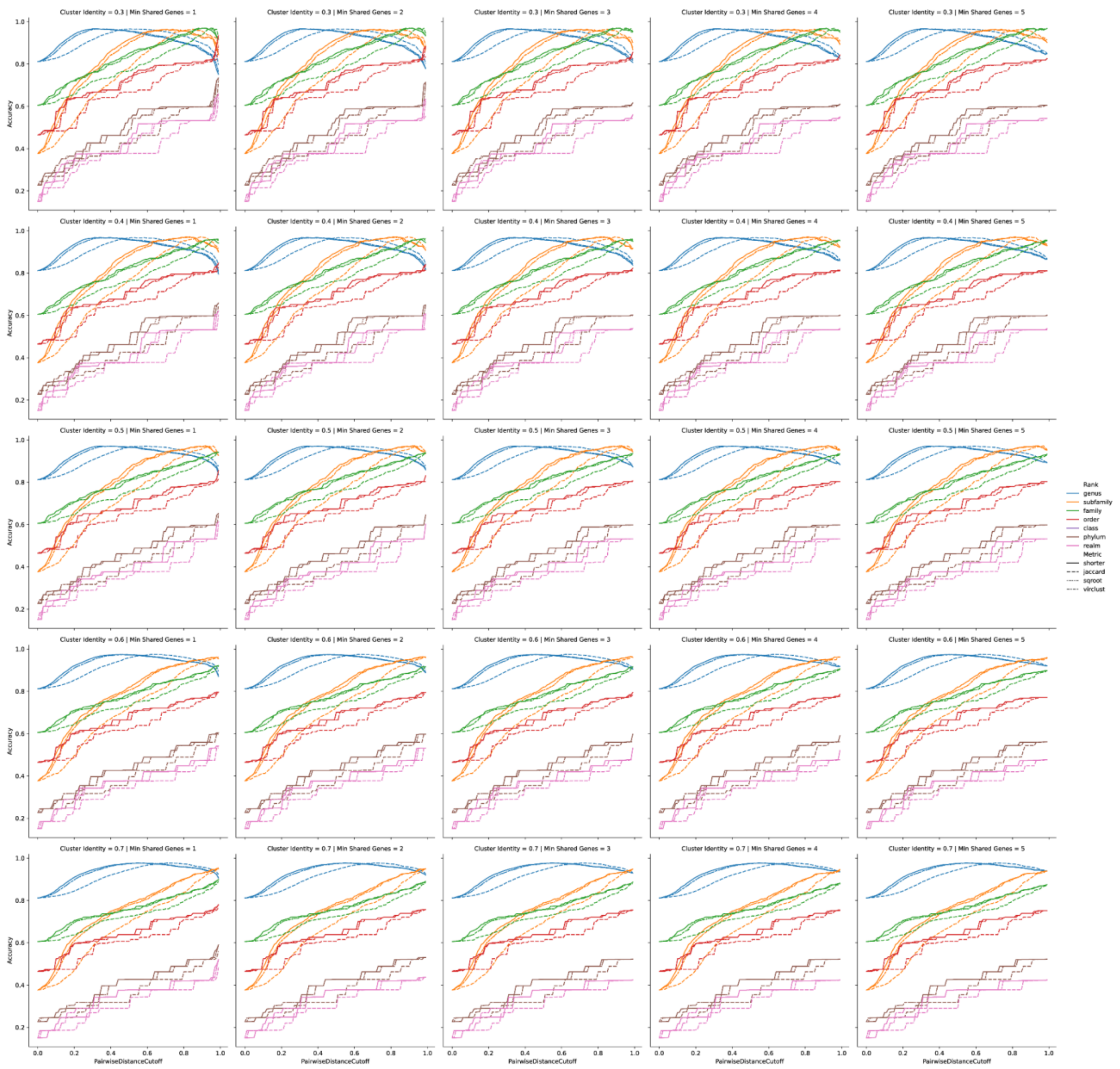


**Extended Data Fig. 6. Prokaryotic-infecting viruses of the *Monodnaviria* realm**. A 5x5 grid of line plots representing accuracies of *Monodnaviria* viruses infecting prokaryotes (*Bacteria* and *Archaea*). Details are as Extended Data Fig. 2.

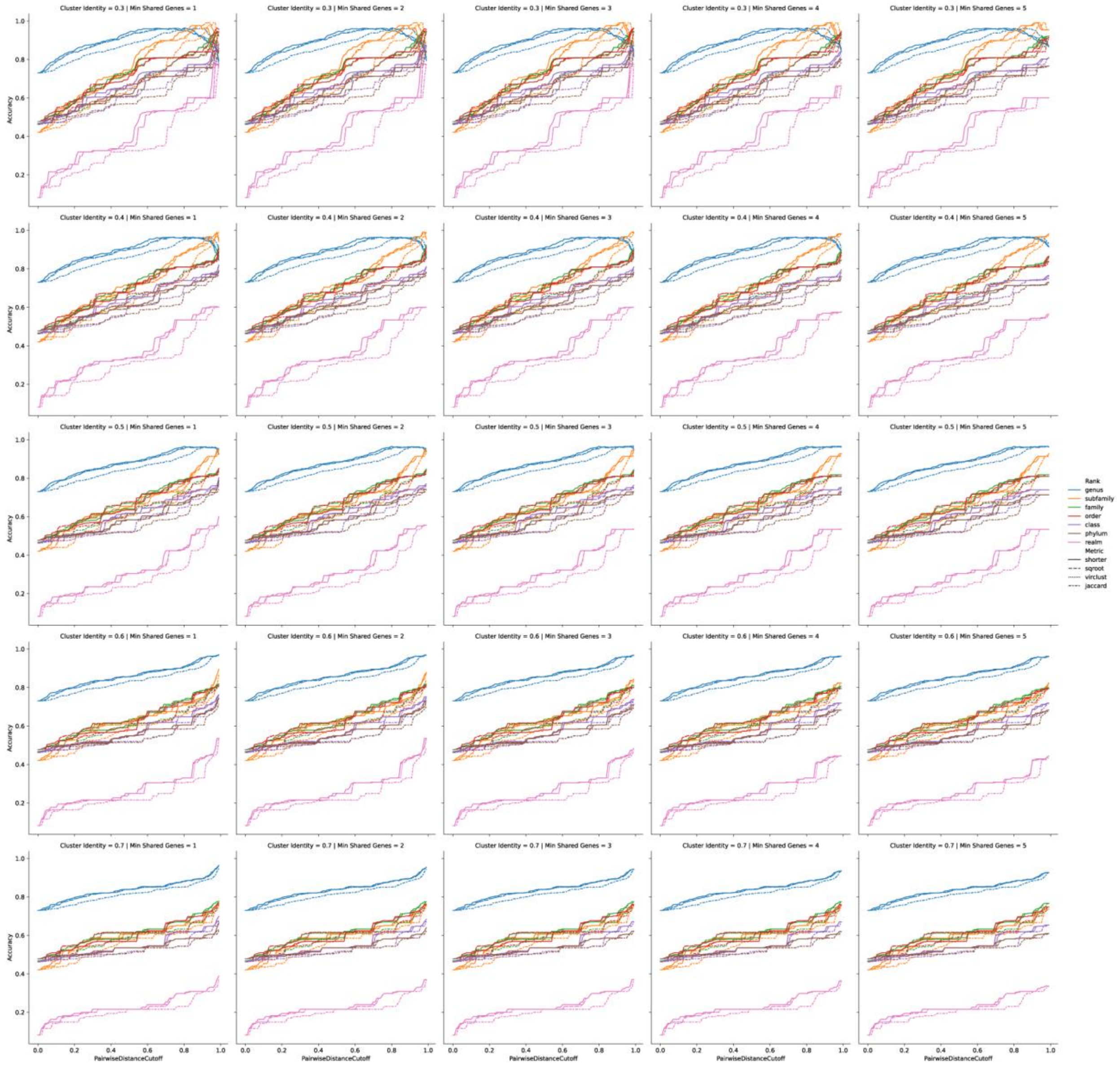


**Extended Data Fig. 7. Eukaryotic-infecting viruses of the *Monodnaviria* realm**. A 5x5 grid of line plots representing accuracies of *Monodnaviria* viruses infecting *Eukaryota*. Details are as Extended Data Fig. 2.

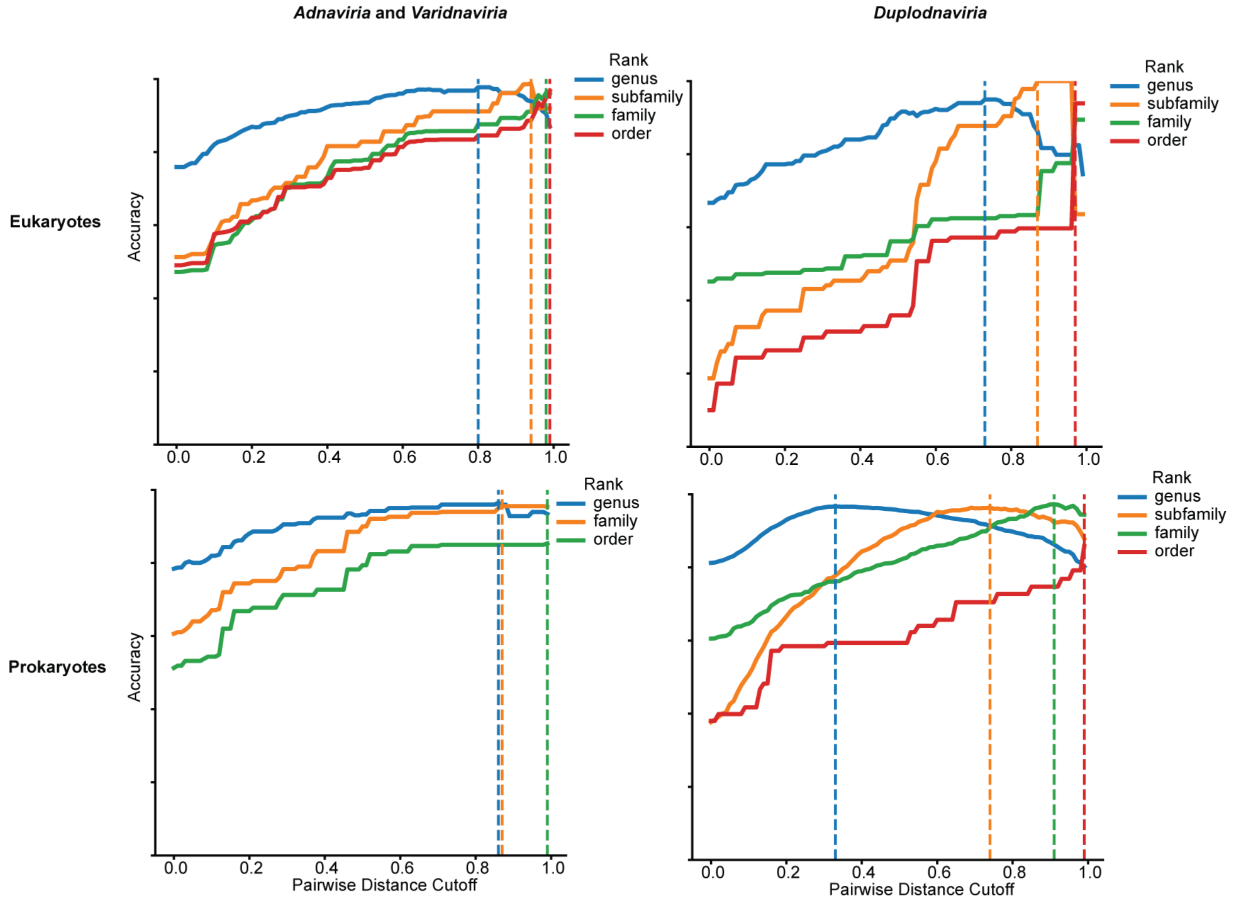


**Extended Data Fig 8. Comparison in distance cutoffs of *Duplodnaviria* and *Adnaviria* & *Varidnaviria* between domains**. Accuracy plots showing the similarity in optimal cutoffs between realms, with prokaryote-infecting viruses appearing downshifted relative to Eukaryote-infecting viruses.

| | genus | subfamily | family | order | class | phylum |
|---|---|---|---|---|---|---|
| *Varidnaviria* | 95.2% | 94.6% | 96.6% | 98.0% | 63.9% | 75.2% |
| *Monodnaviria* | 96.3% | 93.1% | 100.0% | 100.0% | 68.2% | 81.8% |
| *Riboviria* | NP | NP | NP | NP | 41.8% | 54.3% |
| *Duplodnaviria* | 96.5% | 100.0% | 93.7% | 98.3% | 96.4% | 96.4% |

**Extended Data Table 1. vConTACT3 accuracies of Eukaryote-infecting viruses**.
