## Supplemental Information for "Scalable and systematic hierarchical virus taxonomy with vConTACT3"

**Running Title**: Multi-rank, large-scale virus classification with vConTACT3

***Supplemental Note 1: Additional user-friendly outputs new to vConTACT3***

To facilitate virosphere exploration, we improved output files reported to users in several ways. Previously, vConTACT2 provided an assignments table that included references used, some genome statistics, and taxonomic predictions, and also included user-requested outputs increasingly made available in intermediate files. To update this, vConTACT3 now provides outputs as follows. The assignments table now includes more reference information as well as general genome statistics, and taxonomic predictions include suggested “refinement” of existing reference taxonomy where justified. Additional outputs include: (i) a Cytoscape-formatted network with reference and user sequences, predictions, and edge annotations such as distances, weights, and number of genes shared are pre-computed, along with an interactive file (‘d3.js’) that allows users to dynamically explore a network within an HTML interface; (ii) UpSet plots for validating and comparing number and magnitude of edges between and within realms of distance, genome lengths, and shared protein clusters (PCs); (iii) Newick-formatted dendrograms of each connected component in the network to explore results through a phylogenetic-inspired approach; (iv) PC profiles of each user-specified rank for users to identify clade-specific genes for novel virus groups and/or hallmark genes present across multiple ranks; and (v) a completeness table, with estimates based on core genes of references, or a 95% CI using the genus-level group’s sequences when no references exist.

***Supplemental Note 2: Sole exception to cross-realm connections***

There was one exception where viruses from two different realms remained connected, with a link between the *Adnaviria* and *Varidnaviria* realms where 3 shared genes out of 483,002 edges connected 4 genomes (*Sulfolobus islandicus* rod-shaped virus 1 and 2, *Sulfolobus islandicus* rudivirus 3, and *Sulfolobus* turreted icosahedral virus 1). While a cross-realm connection, we consider it valid as the edges shared appear to be host-relevant genes (either involved in CRISPR-Cas defense system or virus egress) likely critical across these dramatically different viruses because they needed similar host-related functions for infection in this extreme environment. Notably, other hosts have extensive virus collections, such as cyanobacteria[33], and these also have shared “host” gene signatures that lead to connections within vConTACT3, but they are less problematic since the virus genomes are much larger and all exist within the same realm. Thus rare multi-realm LCCs may represent critical virus functions or cellular functions.

**Supplemental Note 3: Accuracy and Other Performance Benchmark Comparisons with other Tools**

To measure sensitivity and accuracy of each tool using a set ofwhen compared to known references from, we used the NCBI Virus RefSeq database (release 220). Considering both VPF-Class and geNomad are built on marker genes derived from RefSeq, and vConTACT3 distance thresholds established using RefSeq, we anticipated high rates for all tools. As expected, nearly all tools perform well across the 5 realms, with vConTACT3 and geNomad effectively 100% congruent for phylum and class of Duplodnaviria, Riboviria, and Adnaviria, and order for Adnaviria. geNomad was also fully congruent with Riboviria, whereas vConTACT3 - while capable of identifying RNA viruses - does not classify them beyond class. At order, vConTACT3 was fully congruent with the Duplornaviria, though geNomad suffered a slight drop in accuracy (93%), precision (92%), and sensitivity (93%). At family, both tools performed well, with families under Duplodnaviria and Adnaviria in full agreement, and geNomad achieved 95% accuracy, and 91% accuracy, respectively. The only “available” subfamily was vConTACT3 and Duplodnaviria, which achieved 92% accuracy, 87.5% precision, and 98% sensitivity. Examining genus, where both vConTACT2 and VPF-class are available, vConTACT3 achieved 98.4% accuracy, vConTACT2 96.5%, and VPF-Class 80.8% accuracy within Duplodnaviria. vConTACT3 also achieved the highest levels of sensitivity (99.6%) and precision (97.3%), compared to vConTAC2 (99.9%, 93.1%) and VPF-Class (98.1%, 66.5%). For Adnaviria, both vConTACT3 and VPF-Class performed identically, with 93.5% accuracy, 87.5% precision, and 100% sensitivity. While vConTACT2 had a higher accuracy (96.5%), precision (93%) and sensitivity (100%) versus vConTACT3 and VPF-Class, it also only classified 78% of the sequences, compared to 100% for the other tools. Both vConTACT2 and VPF-Class had 91.2% accuracy, 83% precision, and 100% sensitivity for Riboviria, though only for 4 genomes. For the realm Varidnaviria, both vConTACT3 and geNomad perform similarly, with the former ~4% more accurate at the phylum (78.4%), class (78.4%) and order (100%) ranks. At family, vConTACT3 managed to maintain 100% accuracy, sensitivity, and precision. However, given the low number of genomes included in the analysis (13), we suspect - as with Riboviria - these results may not be representative of the larger virus community. At the genus level, both vConTACT3 and vConTACT2 were 100% accurate, whereas VPF-Class was 90%. As with Adnaviria, vConTACT2 was only able to classify 61% of the realm. While Monodnaviria is not fully optimized with vConTACT3, we still sought to assess its upper rank performance. At phylum and class, geNomad performs best, achieving 99.7% accuracy compared to vConTACT3’s 78.0%. At genus, VPF-Class 91.9% accuracy, 87% precision and 96.9% sensitivity, whereas vConTACT2 performs slightly better, at 93.5%, 88.0% and 99.4%, respectively. We suspect continued optimization of vConTACT3 for this realm will achieve similar or superior results.
